# Stereotyped B-cell responses are linked to IgG constant region polymorphisms in multiple sclerosis

**DOI:** 10.1101/2021.04.23.441098

**Authors:** Ida Lindeman, Justyna Polak, Shuo-Wang Qiao, Trygve Holmøy, Rune A. Høglund, Frode Vartdal, Pål Berg-Hansen, Ludvig M. Sollid, Andreas Lossius

## Abstract

Clonally related B cells infiltrate the brain, meninges and cerebrospinal fluid (CSF) of multiple sclerosis (MS) patients, but the mechanisms driving the B-cell response and shaping the immunoglobulin repertoires remain unclear. Here, we used single-cell full-length RNA-seq and B-cell receptor reconstruction to simultaneously assess the phenotypes, isotypes, constant region polymorphisms, and the paired heavy- and light-chain repertoires in intrathecal B-lineage cells. We detected extensive clonal connections between the memory B cell and antibody-secreting cell (ASC) compartments and observed clonally related cells of different isotypes, including IgM/IgG1, IgG1/IgA1, IgG1/IgG2, and IgM/IgA1. There was a strong dominance of the G1m1 allotype constant region polymorphisms in ASCs, but not in memory B cells. Tightly linked to the G1m1 allotype, we found a preferential pairing of the *IGHV4* gene family with the κ variable *(IGKV)1* gene family. These results link IgG constant region polymorphisms to stereotyped B-cell responses in MS, indicating that the intrathecal B-cell response in these patients could be directed against structurally similar epitopes. The data also suggest that the dominance of the G1m1 allotype in ASCs may occur as a result of biased differentiation of intrathecal memory B cells.

## Introduction

More than half a century ago, Kabat and colleagues discovered that multiple sclerosis (MS) patients have increased levels of immunoglobulin (Ig)G in the cerebrospinal fluid (CSF) (1). Using more sensitive techniques, such as agarose electrophoresis and isoelectric focusing, intrathecally synthesized IgG can be detected as oligoclonal bands (OCBs) in more than 90% of patients (2). OCBs of other isotypes, such as IgM and IgA, can also be found to a variable extent and might be of prognostic value (3–5). The intrathecally produced Igs might originate from long-lived plasma cells residing in survival niches in the central nervous system (CNS) (6) or from short-lived plasmablasts that are being continuously replenished from a pool of memory B cells (7, 8).

A proportion of the secreted IgG proteome in the CSF matches IgG transcripts from B-lineage cells in the brain parenchyma, meninges and the CSF suggesting that CSF IgG is secreted by B-lineage cells within these compartments (9, 10). Accordingly, in the CSF of MS patients, there is an enrichment of antibody-secreting cells (ASCs) that are clonally expanded (11–13), have undergone somatic hypermutation (SHM) (11, 13, 14), and have B-cell receptors (BCRs) displaying a biased usage of variable heavy-chain (V_H_, *IGHV*) genes toward the *IGHV4* family (15). We recently demonstrated that ASCs in the CSF of MS patients are primarily plasmablasts of the IgG1 isotype (8). Surprisingly, in patients carrying the G1m1 and G1m3 allotypes of IgG1, we observed an accumulation of plasmablasts expressing the G1m1 allotype. The mechanism driving this highly selective enrichment of G1m1-expressing ASCs to the CNS is currently not known.

Recent studies have used the 10X Genomics single-cell RNA-sequencing (scRNA-seq) pipeline to study mononuclear cells in the CSF and peripheral blood in MS (13, 16). These studies demonstrated intrathecal T follicular helper cells (16) and clonally expanded B cells (13) with distinct pro-inflammatory profiles compared to their counterparts in blood. Here, we performed an in-depth study of intrathecal B-lineage cells in 21 MS patients using a full-length scRNA-seq protocol (17). We combined this with BraCeR (18), a bioinformatic pipeline that provides reconstruction of paired BCR sequences, clonality inference and lineage tracing of single B cells. This approach has recently proven useful in the study of autoimmune plasma cells in celiac disease (19). In the present study it allowed us to simultaneously assess the detailed phenotypes, isotypes, allotypes, and the full-length paired heavy- and light-chain repertoire of B-lineage cells in the CSF of MS patients.

## Results

### Full-length RNA-seq of single B-lineage cells in the CSF of MS patients

We performed flow cytometry index-sorting and scRNA-seq of B-lineage cells collected from MS patients during diagnostic lumbar puncture (Figure 1, Table 1 and Supplemental Table 1). ASCs were sorted first, and other B-cell phenotypes were included if the number of cells was sufficient (Supplemental Figure 1). After quality control, we analyzed the transcriptomic profiles of 1621 ASCs from 21 MS patients and another 544 B-lineage cells from ten of the patients (Supplemental Tables 2 and 3). All B-lineage cells were phenotyped according to the surface expression of CD19, CD27 and CD38, isotypes, the proportion of Ig transcripts, and the somatic mutation load (Table 2). Based on this, we defined populations of ASCs, naïve and memory B cells. Among CD19^+^ naïve and memory B cells, CD27^+^ memory cells dominated in all patients (median 81.2% [range 66.7-91.4%]; Figure 2A). We further separated memory B cells into non-switched CD27^-/dim^ (4.5% [0-13.3%] of memory cells), non-switched CD27^+^ (36.9% [25.8-60.0%]), switched CD27^-/dim^ (4.4% [0-6.9%]) and switched CD27^+^ memory B cells (54.9% [26.7-65.2%]). Naïve cells constituted a minor part of the sorted B cells in most subjects (8.5% [2.1-34.6%]) (Figure 2A, Supplemental Table 3).

**Figure 1.**
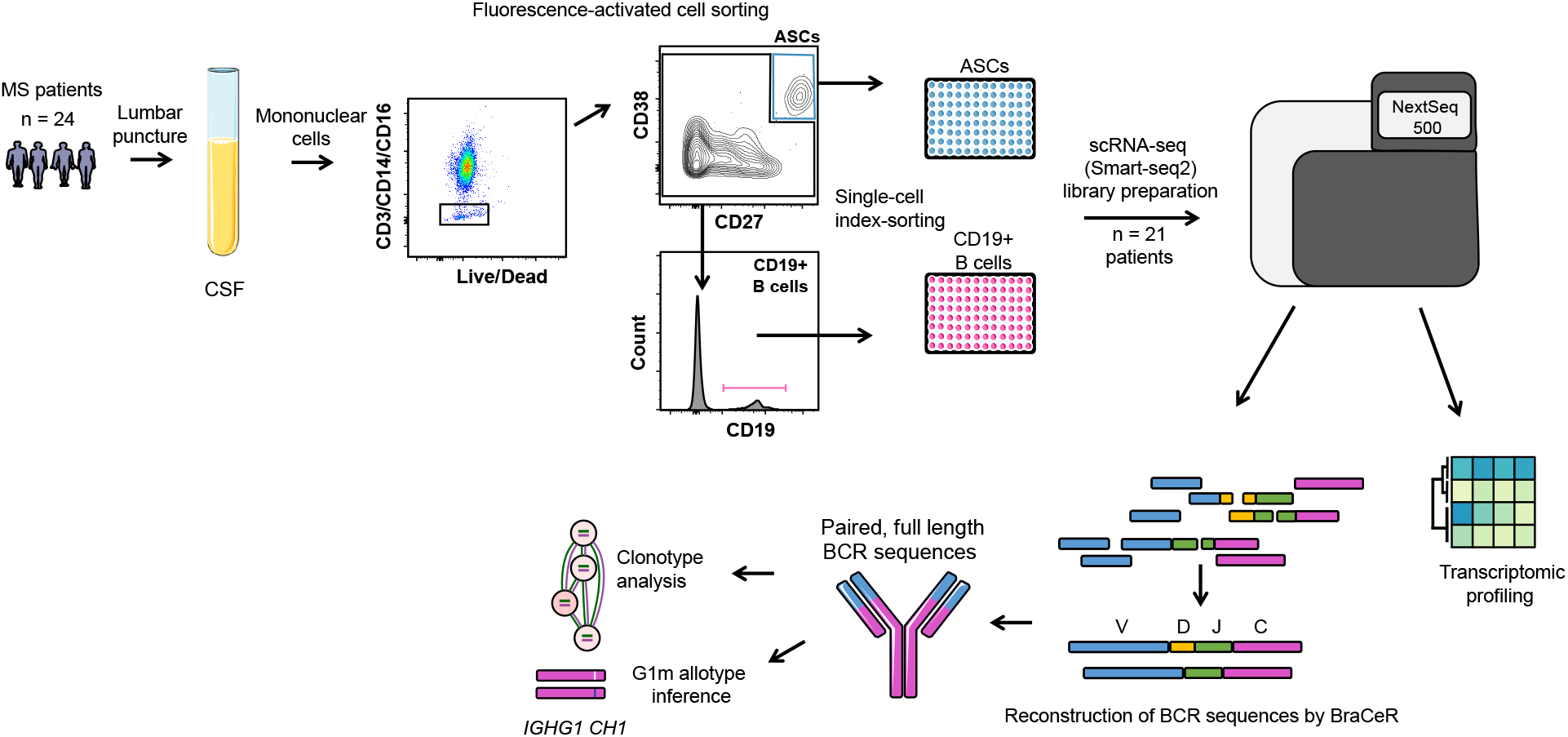
Schematics of single B-cell sorting, sequencing library preparation and data analysis. Fresh CSF samples were stained and analyzed by flow cytometry. Debris and doublets were excluded (not shown). After excluding CD3^+^, CD14^+^ and CD16^+^ cells in a dump channel, CD38++CD27++ single antibody-secreting cells (ASCs) were sorted into 96-well plates containing lysis buffer. For every other plate, we inverted the latter gate and collected single CD38^-/dim^ CD27^-/dim^ CD19^+^ B cells. From the lysed cells, single-cell RNA-sequencing (scRNA-seq) libraries were prepared following a modified Smart-Seq2 protocol and sequenced on an Illumina NextSeq500 platform. B-cell receptor (BCR) sequences were reconstructed using BraCeR. Some of the elements in the figure were modified from Servier Medical Art, licensed under a Creative Common Attribution 3.0 Generic License. http://smart.servier.com.

**Figure 2.**
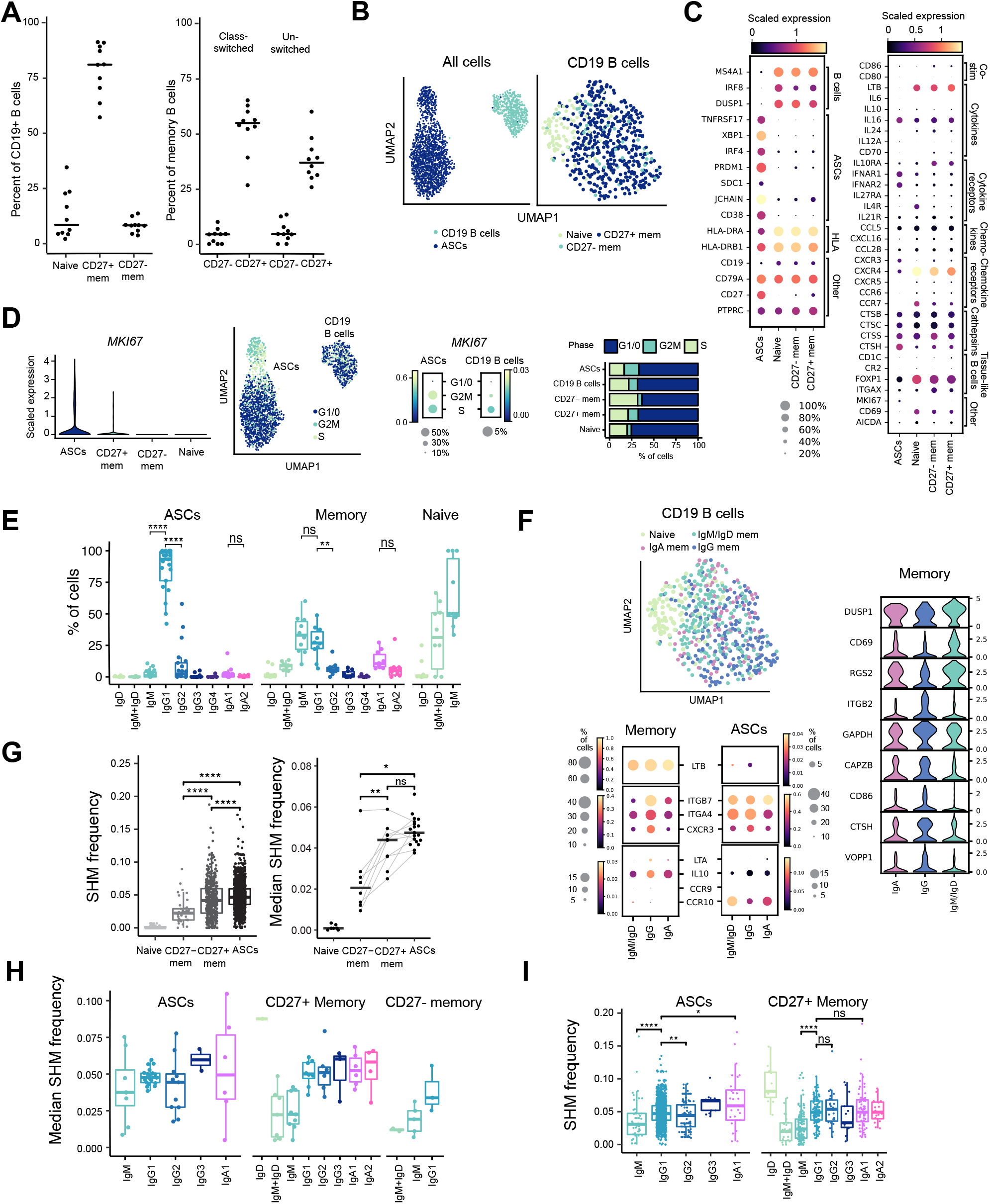
Transcriptional and mutational profiling of B-lineage cells from the cerebrospinal fluid of MS patients. **A.** Frequencies of sorted CD19^+^ B cells being classified as naïve or memory B-cell subsets (Table 2). **B.** UMAP projection of all B-lineage cells (left; n=21, 2165 cells) or only the CD19^+^ B cells (right; n=10, 544 cells). **C.** Heatmap showing expression of genes of particular interest in all B-lineage cells in (B). ASC: antibody-secreting cell, HLA: human leukocyte antigen. **D.** *MKI67* expression and inferred cell cycle phase in each B-lineage population. **E.** Isotype distribution based on reconstructed B-cell receptor (BCR) sequences. Each data point represents a patient with more than one cell for a given cell type. **F.** Differentially expressed genes (DEGs) between B-lineage cells of different isotypes. The UMAP of CD19^+^ B cells (upper left) is identical to (B), but colors refer to naïve cells and the isotype of memory cells. For gene expression analysis of memory B cells, naïve B cells were removed from the scaled scRNA-seq expression data for CD19^+^ B cells, and the cells were grouped according to isotype (198 IgM/IgD, 172 IgG, 86 IgA cells). Gene expression analysis was performed separately for the ASCs (55 IgM/IgD, 1529 IgG, 34 IgA cells). Statistically significant DEGs for the memory B cells (Wilcoxon rank-sum test adjusted for multiple testing) with a log fold change greater than 1 are displayed in a violin plot (right). **G.** Somatic hypermutation (SHM) frequency in the heavy- and light-chain variable region of B-lineage cells by phenotype. Left panel shows all BCR sequences (Wilcoxon rank-sum test); right panel shows median frequency for each patient (Wilcoxon signed-rank test, excluding unpaired data points). Grey lines connect data points from the same patient for memory B cells and ASCs. **H,I.** SHM frequency in the heavy- and light-chain variable region of B-lineage cells by phenotype and isotype. Median frequency for each patient (H) or frequency in all BCR sequences (I; Wilcoxon ranksum test) are shown. P-values throughout Figure 2 are given with the following significance levels: *: p<0.05; **: p<0.01; ****: p<0.0001, ns: p>0.05.

**Table 1.**
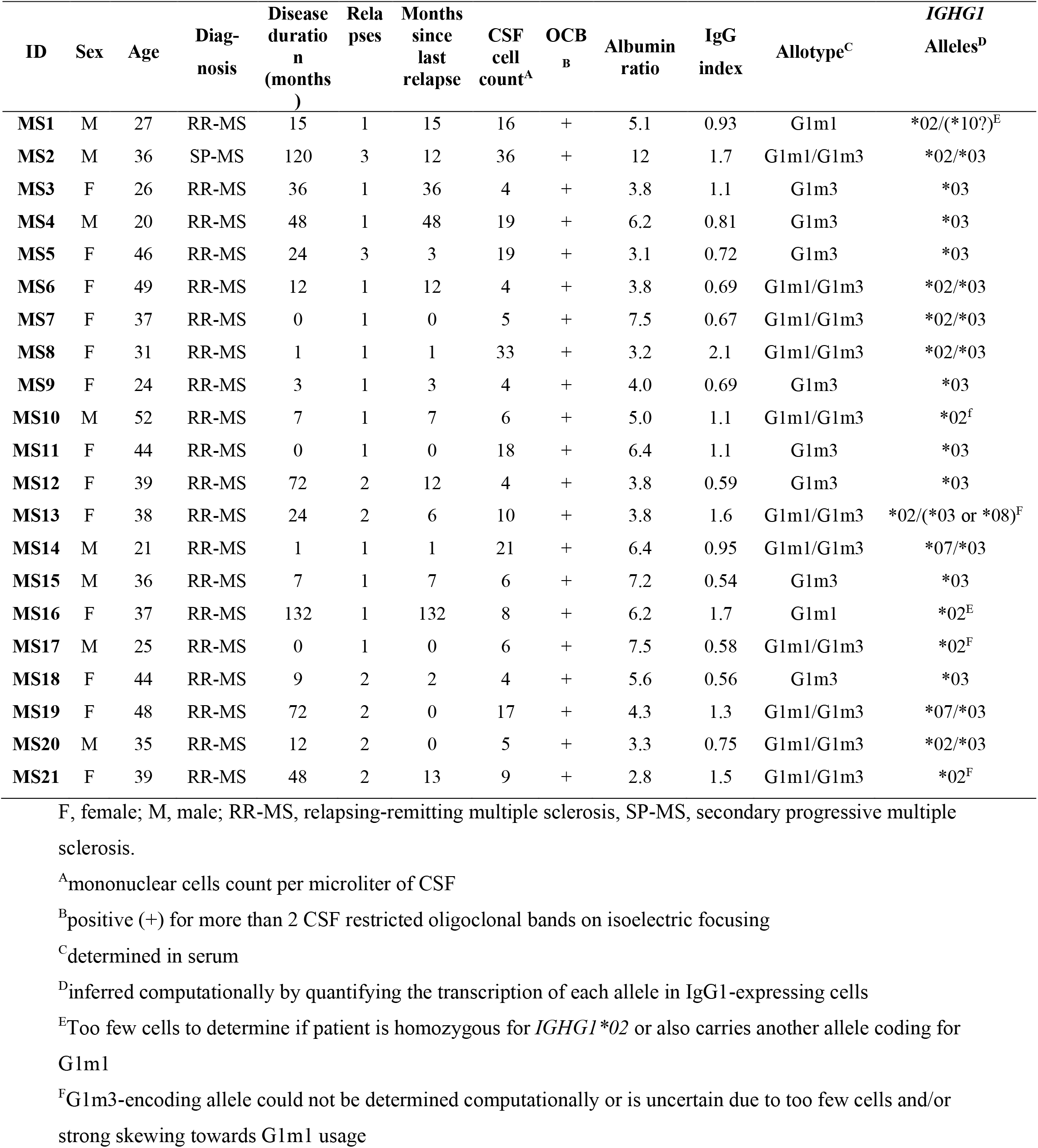
Patient characteristics.

**Table 2.**
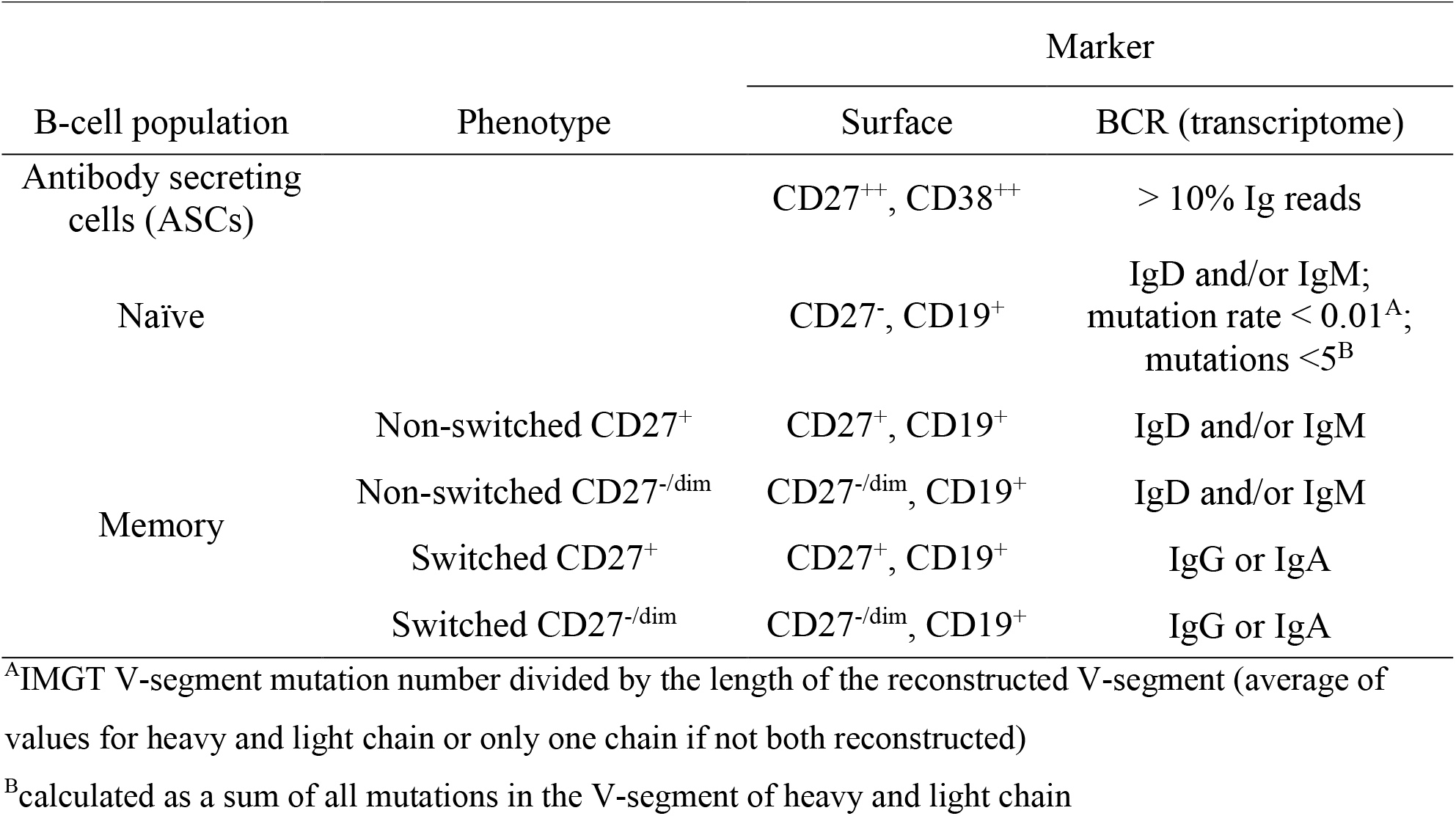
Features used to phenotype B-lineage cells.

### B-lineage cells in the CSF of MS patients express genes involved in B-cell differentiation, antigen presentation and cytokine production and signaling

The transcriptomes of these cell populations were visualized using UMAP (Figure 2B and Supplemental Figure 2, A and B). As expected, ASCs clustered separately from the CD19^+^ B cells. Within the CD19^+^ B-cell population, naïve B cells clustered separately from memory B cells. Next, we analyzed the expression of genes in the main B-lineage populations (Figure 2C). The ASCs expressed typical plasmablast/plasma cell genes such as *XBP1, IRF4, PRDM1* (BLIMP-1), and a proportion (45%) expressed *SDC1* (CD138). We also searched for expression of genes associated with antigen presentation and detected ubiquitous expression of human leukocyte antigen (HLA) class II genes, in addition to cathepsins (Figure 2C). Next, we looked at cytokine and chemokine expression. A proportion (19%) of the memory B cells showed expression of *CD70* (encoding the ligand for CD27), which was absent in the naïve B cells and barely detectable in the ASCs. Further, most memory and naïve B cells expressed high levels of *LTB*, whereas *IL16* and to a much lower degree *IL10* was expressed in a proportion of all cell subsets. Chemokine expression was detected to some extent, however we found broad expression of genes encoding cytokine/chemokine receptors in all populations. Memory and naïve populations expressed high levels of *CXCR4*, in addition to restricted expression of *CXCR3, CXCR5, CCR6, CCR7* and *IL10RA* among others, while ASCs expressed genes such as *TNFRSF17, IFNAR2* and *IL21R* and also *CXCR3* and *CXCR4* (Figure 2C).

### ASCs in the CSF constitute a continuity from immature plasmablasts to more differentiated phenotypes

Approximately 22% of the ASCs expressed the proliferation marker *MKI67* (Ki-67), suggesting they were plasmablasts rather than end-differentiated plasma cells (Figure 2D and Supplemental Figure 2C). Notably, the ASCs clustered based on inferred cell cycle phase when visualized by UMAP (Figure 2D), and *MKI67* was as anticipated expressed in cells designated to the G2M or S phases. This held true also when excluding *MKI67* from the list of genes used to infer cell cycle phase (Supplemental Figure 2C). When visualized by UMAP, ASCs assigned to the non-proliferative G1/0 phase (66.9%) displayed a seemingly gradual transition between a major population of ASCs with high expression of *SDC1* (CD138) and a smaller population expressing *MS4A1* (CD20) (Supplemental Figure 2D). These data suggest that ASCs in the CSF of MS patients are proliferating plasmablasts within different stages of maturation. However, a proportion of the ASCs in the inferred G1/0 phase may represent true end-differentiated non-dividing plasma cells.

### IgG, IgA and IgM/IgD B-lineage cells show distinct transcriptional profiles and mutational load

Full-length recombined BCR sequences, stretching into the constant region, allowed us to define the isotype of sorted B-lineage cells. In line with previous studies (8, 20), IgG1 was by far the most prevalent isotype among ASCs (Figure 2E). The median proportion of ASCs expressing the IgG1 isotype was 93.3% [42.1-100%], while in the memory subset the IgM and IgG1 isotypes were used the most (33.0% [10.0-60.0%] and 26.9% [6.7-48.8%], respectively). In line with previous reports of isotype-specific transcriptional profiles of B-lineage cells (21), we observed a gradual transition between naïve, non-class-switched and class-switched memory B cells when visualized by UMAP (Figure 2F). Genes that were found to be significantly higher expressed in memory B cells of the IgM/IgD compared to IgG and IgA isotypes included *DUSP1, CD69*, and *RGS2* (encoding Regulator of G-protein signaling 2). *CTSH* (encoding Cathepsin H) and *CD86* were significantly higher expressed in IgG memory B cells, suggesting an increased importance of antigen presentation in these cells. IgG memory B cells also had the highest expression of CXCR3, which is shown to be important for recruitment of B-lineage cells to inflamed tissues (22). In contrast to a recent report (23), we did not detect a distinct regulatory phenotype of IgA-expressing B-lineage cells in the CSF, and we did not find evidence to support that IgA-expressing B-lineage cells express higher levels of gut-homing markers than their IgM/IgD and IgG counterparts (Figure 2F).

Next, we investigated the mutational load of each reconstructed BCR sequence. Numbers of somatic mutations varied between populations (Figure 2G) and isotype (sub)classes (Figure 2, H and I). In line with previous reports (24, 25), CD27^-/dim^ memory B cells displayed significantly fewer mutations than their CD27^+^ counterparts. Among ASCs we observed the highest number of mutations in class-switched cells. In the memory B cell population, IgD-only cells showed the highest mutational load, followed by class-switched memory cells (Figure 2, H and I).

### B-lineage cells in the CSF of MS patients undergo intrathecal maturation

Next, we analyzed clonal relationships between B-lineage cells using BraCeR. BraCeR, for paired heavy and light chains, assigns clonal relationships between cells, which are then depicted as clonal networks and additionally lineage trees for individual clones consisting of cells with different mutational patterns (Figure 3). In all 21 patients clonally related cells were detected, with a median of 61.6% [39.7-83.1%] of all ASCs belonging to a clone (Figure 3, A and B, and Supplemental Table 4). As the most extensive study to date, we traced the clonal evolution of B-lineage cells based on somatic mutations in the paired full-length variable heavy- and light-chain genes. Thus, we reconstructed lineage trees in 17/21 patients (average 2.4 unique sequences per lineage tree [range 2-8]; Figure 3C and Supplemental Table 4). Interestingly, we commonly observed highly mutated clones consisting solely of cells with identical mutational patterns (Supplemental Table 4). The four patients with no reconstructed lineage trees had low numbers of ASCs and/or the sequences had the exact same mutational patterns.

**Figure 3.**
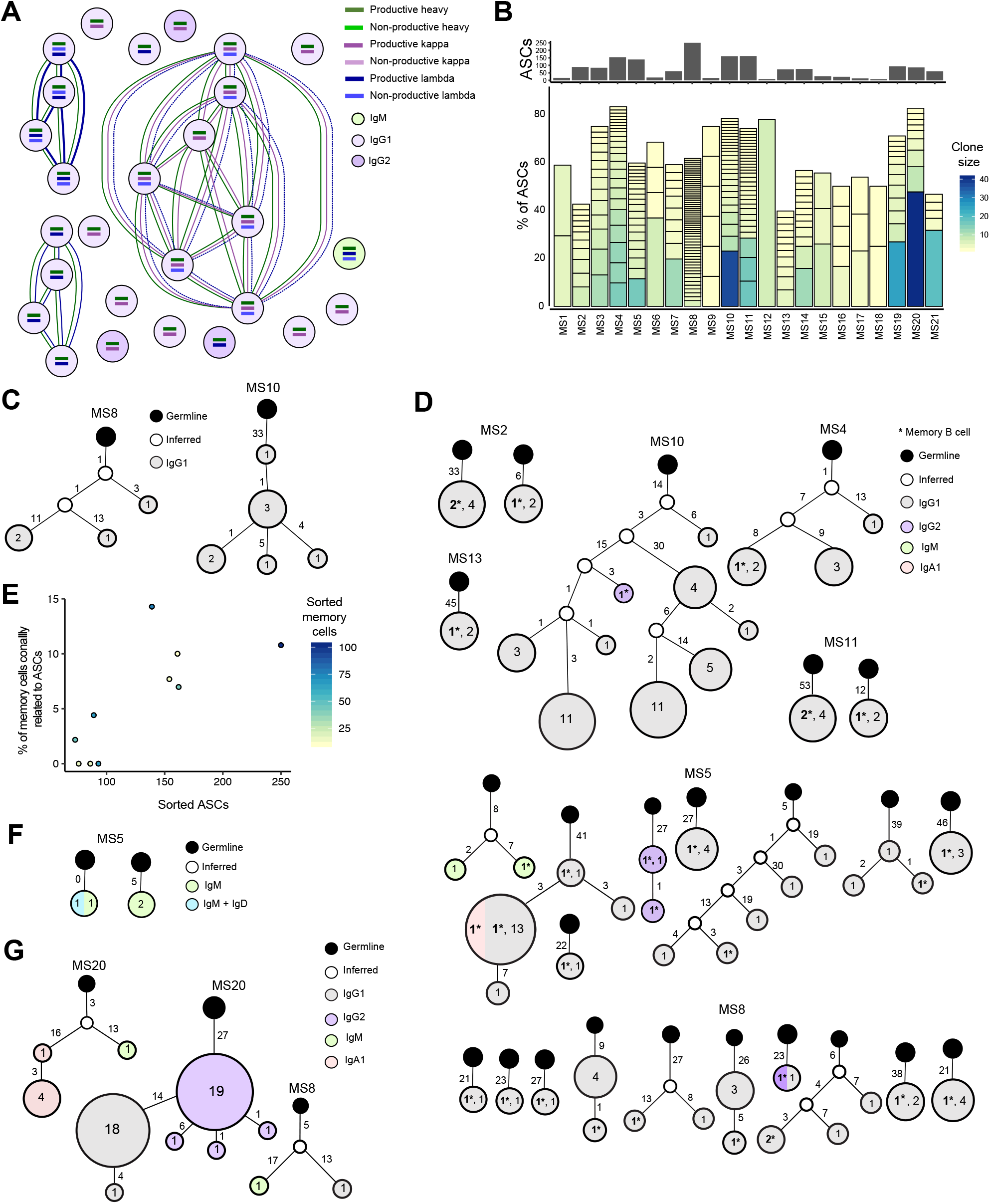
B-lineage cells in the CSF are clonally related, undergo somatic hypermutation, isotype switching and intrathecal maturation. **A.** Example of clonal relationships between antibody secreting cells (ASCs) of patient MS15 inferred by BraCeR. **B**. Sizes of clonal groups found for ASCs in each patient. The colors indicate number of cells belonging to each clone group, size of boxes is proportionate to the size of clones. Only clonally expanded cells are shown. **C.** Representative ASC lineage trees generated by BraCeR for two clones, with inferred germinal sequences at the root. The number of mutations between each node is shown next to the branch. The size of each node is proportionate to the number of cells containing a given unique BCR sequence. The number inside each node represents number of cells with the unique sequence. **D.** Clones comprising both memory B cells and ASCs, visualized as lineage trees as in (C). **E.** Relationship between number of sequenced memory B cells, ASCs and percent of memory B cells found to be clonally linked to ASCs. Each dot represents one patient. **F.** Clones limited to the CD19^+^ B cell compartment, visualized as lineage trees. **G.** Examples of multiple isotypes found within B-lineage clones, visualized as lineage trees.

Our approach, where both heavy and light chains are involved in determining clonal relations combined with transcriptomic profiling of cells, made it possible to trace linkage between memory B cells and ASCs. Strikingly, we observed clonal sharing between memory B and ASCs in 7 out of 10 patients from whom both cell populations were sorted (Figure 3, D and E). The majority of memory cells belonging to clones containing ASCs shared identical V-regions with at least one ASC, while we also observed acquisition of mutations. In one patient, we found clonally expanded memory B cells with no detected members of the clones among the ASCs (Figure 3F). The lack of connections between memory B cells and ASC in three patients could possibly be attributed to low cell numbers (Figure 3E). We also explored the clonal evolution of the ASCs in relation to their cell cycle stage and found extensive clonal connections between cells in different stages (Supplemental Figure 3, A-D).

The most commonly found expanded sequences other than IgG1 were of the IgG2 isotype subclass (Table 3, Supplemental Figure 3E). In eight patients we also found expanded IgA1 and/or IgM ASCs. Additionally, we detected expanded IgG3 ASCs in two patients and expanded IgG4 ASCs in one patient. Interestingly, we found examples of clones comprising B-lineage cells of different isotypes, including IgM and IgG1, IgG1 and IgA1, IgG1 and IgG2, and IgM and IgA1 (Figure 3, D and G). This could suggest that class-switch recombination has taken place intrathecally and that at least a proportion of the IgG, IgM and IgA B-lineage cells share a common origin and target the same antigens.

**Table 3.** Clonal expansion of other isotvpes than IsG1.

| ID | IgA1 | IgM | IgG2 | IgG3 | IgG4 |
| --- | --- | --- | --- | --- | --- |
| MS2 | - | - | 1(2) | - | - |
| MS3 | - | 1(2) | 1(11) | - | - |
| MS4 | - | 2(3) | 1(4) | - | - |
| MS5 | - | 2(3-5) | 1(2) | 1 (6) | - |
|  | - | 2(2) | 1(2) | - | - |
| MS6 | - | - | 2(2-7) | - | - |
| MS8 | 1(2) | - | - | - | - |
| MS9 | 1(2) | - | 1(2) | - | - |
| MS10 | 2 (2) | - | - | - | - |
| MS11 | 1 (2) | 1(3) | 2(2-3) | 1(3) | - |
| MS14 | - | - | 2(7-12) | - | 1(2) |
| MS20 | 1(5<sup>A</sup>) | 1(2) | 3(2-22<sup>B</sup>) | - | - |

Numbers of clonotypes for each patient with numbers in brackets indicating sizes of clonal expansion. All cells are expanded ASCs except the shaded, which are expanded memory B cells.
<sup>A</sup>Clone additionally contains one IgM ASC.
<sup>B</sup>Part of a large expanded clone containing 22 IgG2 and 19 IgG1 ASCs.

### ASCs but not memory B cells show a strong preferential usage of the G1m1 allotype

BraCeR infers the CH1 region of the *IGHC* alleles of reconstructed heavy chains, allowing for detection of isotypes and distinction between the G1m1 and G1m3 allotypes on a cellular level. Due to random allelic exclusion of Ig genes, there is a 50/50 distribution of the *IGHC* alleles among B-lineage cells in the blood of heterozygous individuals (8). We have previously shown, based on flow cytometry, that G1m1-expressing ASCs are preferentially enriched in the CSF of MS patients (8). To address this on a transcriptomic level, we limited the analysis to G1m1/G1m3 heterozygous patients and calculated the proportion of G1m1-expressing cells among IgG1 ASCs and memory B cells (Figure 4, A and B). In agreement with our previous study, the results showed a predominance of G1m1 ASCs (Figure 4A; median 97.8% [66.7-100%]). On an individual level, a statistically significant skewing towards the G1m1 allotype in the ASC population was seen in 10/11 of the G1m1/G1m3 heterozygous patients (Figure 4B). In contrast, the G1m1 bias was not present to the same degree in the IgG1 memory B cell population (Figure 4, A and B).

**Figure 4.**
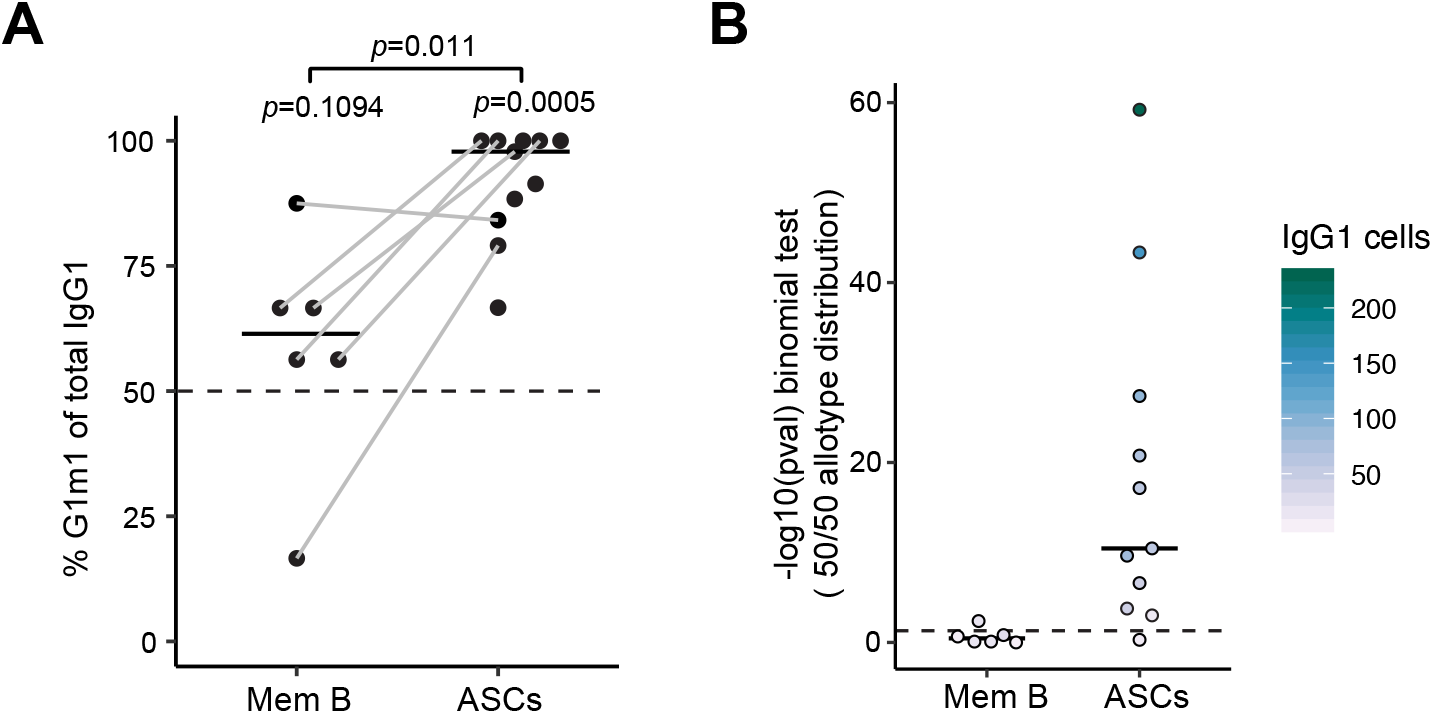
The intrathecal ASC population is skewed toward the G1m1 allotype in G1m1/G1m3 heterozygous MS patients. **A.** G1m1 allotype distribution in IgG1 memory B cells (n=6) and antibody-secreting cells (ASCs; n=11) of G1m1/G1m3 heterozygous patients. Each dot represents the percentage of IgG1 cells that are of the G1m1 allotype for each patient and horizontal solid lines depict median for all patients. Lines connect ASCs and memory B cells from the same patients. Frequencies of G1m1 cells were compared within each population (one-sided binomial test) and between paired memory and ASCs populations (paired t-test), where four patients were excluded due to absence of a memory cell population (no CD38^dim/-^CD27^dim/-^CD19^+^ cells sorted) and one patient due to only one IgG1 memory B cell being present for this patient. The horizontal dashed line at 50% represents the expected distribution of G1m1 cells due to random allelic exclusion. **B.** P-values resulting from two-sided binomial tests for each population within each patient. Solid horizontal lines indicate median values. The horizontal dashed line represents statistical significance (p=0.05).

### ASCs expressing G1m1 predominantly use IGHV4 gene segments, k light chain and show preferential pairing of V_H_ and V_L_ genes

Previous studies have shown that the *IGHV4* gene segments are dominating the CNS and CSF heavychain repertoires (15, 26). We stratified the heavy-chain repertoires of ASCs according to G1m allotype and found that a preferential use of *IGHV4* gene segments is associated with the G1m1 allotype, but not with the G1m3 allotype (Figure 5A and Supplemental Figure 4, A and B). In the G1m1-expressing memory B-cell population, on the other hand, we did not detect a similar bias of *IGHV4* gene segments usage (Figure 5A). To confirm the association between *IGHV4* gene segments and G1m1 in a completely independent patient cohort, we reanalyzed *IGHV* sequencing data from bulk CSF B-lineage cells of 12 MS patients previously published by us (27, 28). We used total *IGHV* pool, which drives bias toward ASCs-derived sequences (29), and it yielded similar results (Figure 5B). Next, we looked into the repertoire of the light chains in single IgG1 ASCs. The results showed that a strong preference for the κ light chain was connected to the G1m1 allotype, but not the G1m3 allotype (Figure 5C and Supplemental Figure 5A). No significant λ light-chain bias was seen for G1m1-expressing memory B cells, but this could possibly be attributed to few data points and large variation (Figure 5C). We also investigated the use of light-chain constant region genes and alleles (Supplemental Figure 5, B and C). In agreement with the expected high allele frequency of the Km3 allotype in Caucasians (30), we detected use of the Km3-encoding *IGKC*01* allele in all patients with κ-expressing cells. Two patients were heterozygous for *IGKC*01* and the Km1,2-encoding *IGKC*04* alleles, and exhibited no preferential usage of either allele (Supplemental Figure 5C). No serological allotypes are known for the λ light chain, which instead exhibits variation through a variable number of different *IGLC* genes. We observed no strong preference of any specific *IGLC* genes in G1m1 and G1m3 cells (Supplemental Figure 5C).

**Figure 5.**
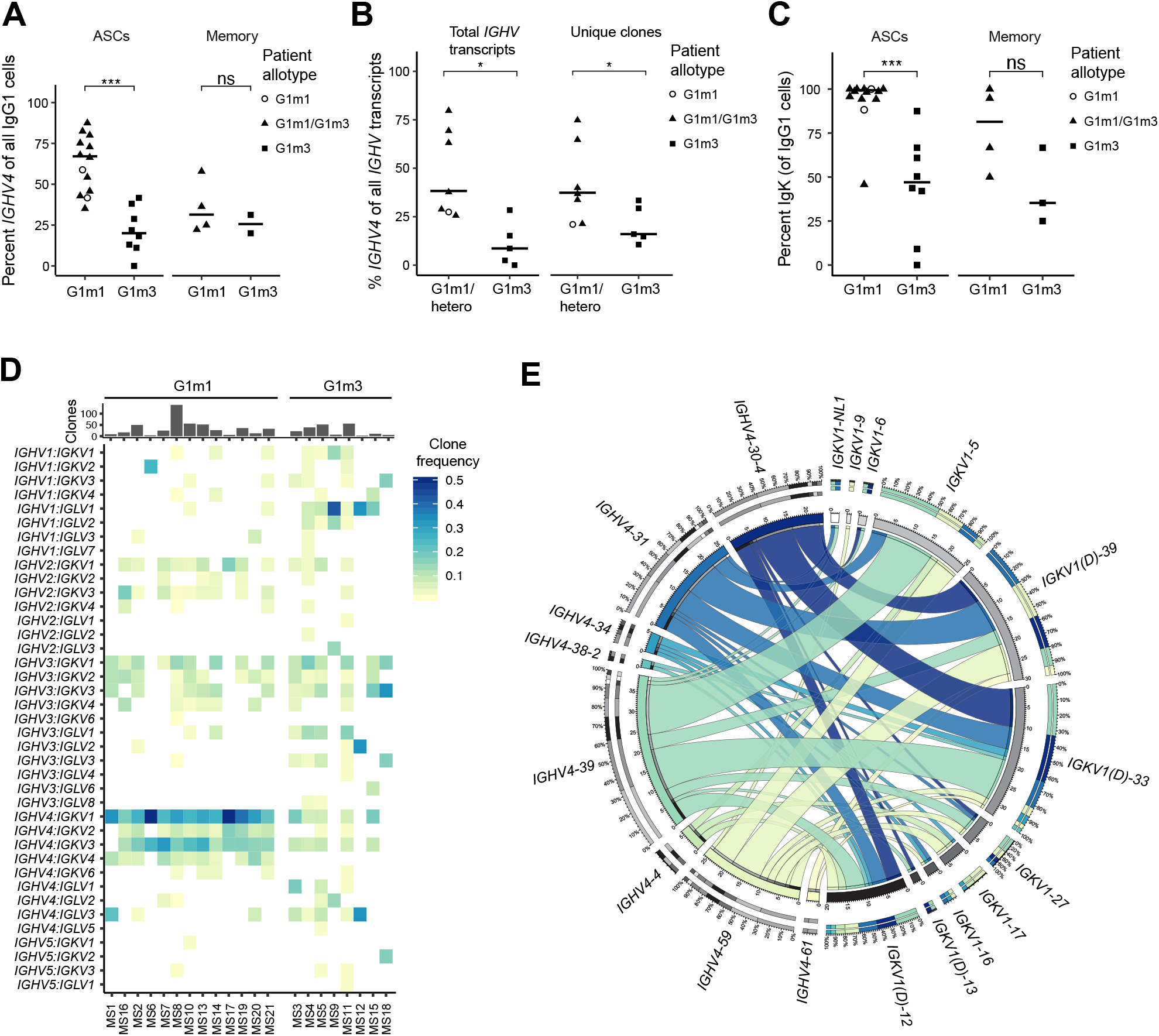
The G1m1 allotype is associated with a stereotyped BCR repertoire. Analysis was performed on the V_H_ and V_L_ sequences of IgG1 antibody-secreting cells (ASCs). G1m1 cells from G1m1 homozygous and G1m1/G1m3 heterozygous patients are considered G1m1, whereas only sequences from G1m3 homozygous individuals are denoted as G1m3. G1m3 cells from heterozygous patients were excluded from the analysis due to low cell numbers. A Wilcoxon ranksum test was used in (A-C), with p-value significance levels *: p<0.05; ***; p<0.001; ns: p>0.05. **A.** Percentage of G1m1 and G1m3 cells using *IGHV4* gene segments. Each dot represents a patient and is shaped according to patient G1m1/G1m3 carrier status; horizontal line represents median value. **B.** A reanalysis of previously published bulk RNA-seq data of *IGHV* genes.(27, 28) Patients were grouped according to presence (n=7) or absence (n=5) of the G1m1 allotype by serotyping. **C.** κ light chain usage of G1m1 and G1m3 ASCs. Each dot represents a patient; horizontal lines represent median values. **D.** Heatmap showing V_H_:V_L_ gene segment pairing frequencies for G1m1 and G1m3 cells, where clonally expanded cells are treated as one cell. **E.** Circos plot visualizing the *IGHV4:IGKV1* gene segment pairings for G1m1 ASCs, where each clone group is treated as one sequence. Pairing is depicted as lines connecting *IGHV4* and *IGKV1* genes inside the circle, where thickness corresponds to pairing frequencies. Sequences from all G1m1 carriers (n=13) were pooled together.

The analysis of single B-lineage cells allowed us to gain insight into the V_H_:V_L_ pairings, which is key to understanding antigen-driven B-cell responses and may sometimes show specific signatory combinations (31, 32). In order to exclude the influence of clonal expansion on the pairing frequencies, only one sequence per clonotype was taken into account. In ASCs expressing G1m1, we observed a preferential pairing of *IGHV4* with *IGKV1* and to a lesser degree *IGKV3* gene segments (Figure 5D). The same pairing preference was not observed for G1m3-expressing ASCs. This pattern was also evident when taking clonal expansion into account (Supplemental Figure 4C), but it was not clear in the naïve and memory B-cell population (Supplemental Figure 4D). Finally, we explored the pairing frequencies of genes within the *IGHV4* and *IGKV1* gene families in G1m1-expressing ASCs. The *IGHV4-39* gene was most commonly used and showed the highest frequency of pairing with *IGKV1-5* (frequency of 9.9%) and *IGKV1(D)-33* (8.4%) (Figure 5E).

## Discussion

In the present study we analyzed the paired full-length rearranged Ig heavy- and light chain gene usage and transcriptome of intrathecal B-lineage cells in MS. We achieved this by combining full-length transcriptomic profiling of single cells along with analysis using BraCeR. The results reveal a common pattern across patients with a preferential V_H_:V_L_ pairing tightly linked to the G1m1 allotype in ASCs. Confirming our previous findings on a transcriptional level (8), the G1m1-expressing ASCs were enriched in the CSF of G1m1/G1m3 heterozygous patients. Our approach further allowed us to demonstrate (i) clonally expanded IgG1, IgG2, IgG3, IgG4, IgA1 and IgM B-lineage cells and clonal connections between them, (ii) accurate lineage trees based on mutations in both heavy- and lightchain genes of clonally expanded ASCs, and (iii) the presence of expanded memory B cells, a few examples of isotype switching and extensive clonal connections between the memory B-cell and ASC compartments.

Previous studies have demonstrated that Ig heavy-chain sequences from the brain, CSF and cervical lymph nodes of MS patients are clonally expanded and have undergone SHM, which is indicative of an antigen-driven immune response (11, 13–15, 27, 33–35). *IGHV4* gene segments have been portrayed as dominant in the CSF and brain of patients with MS and clinically isolated syndrome (15, 26, 36), and have also been shown to acquire specific mutations and displaying increased specificity toward neuroantigens (37, 38). At a closer look, however, it seems that a *IGHV4* bias is only present in a proportion of patients, although this has not been made a matter of contention by previous studies (27, 33). The present results offer a potential explanation, which is possible to verify in previously published data if the G1m1 carrier status of the patients is determined. Indeed, reanalyzing data from our own group that were generated using bulk sequencing of CSF B cells and a multiplex PCR technique confirmed the association between the G1m1 carrier status and a biased usage of *IGHV4* gene segments (27, 28). Curiously, the present data also show a connection between G1m1 expression and λ light-chain usage. While λ light-chain usage was highly dominant among ASCs of G1m1-carriers, several G1m3 homozygous patients showed a predominance of λ light-chains in the ASC population. Thus, the G1m1 carrier status could have implications for the diagnostic sensitivity of free light-chain levels in the CSF, which is increasingly being recognized as a valuable diagnostic test in MS (39) and might correlate with the frequency of ASCs expressing the light chain (40).

Preferential pairing of *IGHV4* and *IGKV1* in the BCR of ASCs expressing G1m1 bears a resemblance to stereotyped B-cell responses observed in other diseases, including celiac disease (41), HIV infection (42), and influenza infection (43) that are driven by particular antigenic epitopes (44). The target antigens of the intrathecal humoral immune response in MS, however, have not been unequivocally defined. Some early studies reported reactivity against myelin-associated antigens (45, 46) and Epstein-Barr virus (47), but none of these findings have been reproduced in independent studies (12, 48). More recent studies have shown that some antibodies expressed by CSF B cells recognize neuronal nuclei and/or astrocytes (49) and lead to demyelination in spinal cord explant cultures (50). Finally, a study reported that a proportion of CSF IgG might target cellular debris (51). Nevertheless, independent of the nature of the antigen, the stereotyped B-cell response with a preferential V_H_:V_L_ pairing demonstrated here argues that these B-lineage cells target a set of epitopes that may be shared between patients.

We have previously demonstrated that the G1m1 allotype dominates the intrathecal humoral immune response in G1m1/G1m3 heterozygous MS patients, but not in controls with neuroborreliosis (8). The present study confirms the observation in MS patients using gene expression analysis of single CSF ASCs and extends the findings demonstrating that the G1m1 bias is not present to the same degree in intrathecal memory B cells. Considering the extensive clonal connections between these intrathecal populations, this could indicate that the dominance of the G1m1 allotype in ASCs is the result of a preferential differentiation of certain B-lineage cells. The mechanisms driving such a biased maturation is currently not known, but one important clue comes from the findings of a connection between the G1m1 allotype and particular V_H_ and V_L_ gene segments, suggesting that the ASCs target certain antigenic epitopes. A genetic linkage between the alleles containing G1m1 and certain *IGHV* genes on chromosome 14 and/or conformational changes in the variable antigen-binding region induced by the G1m allotypes in the constant region are potential explanations to how an antigen-driven B-cell response could show such a preference. In support of the latter idea, it has been shown that the constant region of Igs may affect the affinity and specificity of the variable region (52). The biased usage of *IGKV* gene segments located on chromosome 2, on the other hand, could be the result of interactions with the expressed heavy chain and the antigen. It is conceivable that the distinct BCR repertoire we observe in patients carrying the G1m1 allotype may have consequences for disease risk and phenotype. Accordingly, it is known that genetic markers in linkage disequilibrium with the G1m1-encoding alleles are associated with higher IgG index (53) and lower IgM index (54), and possibly with MS risk (54). Further studies investigating the association with the disease risk and phenotype are warranted.

The last several years, great progress has been made in our understanding of the trafficking of B-lineage cells between the periphery and the CNS (27, 33, 34, 55). Accordingly, at least a proportion of B-lineage cells within the brain have been shown to originate in deep cervical lymph nodes (34). Further, a number of different B-cell populations in blood seem to have connections to the intrathecal B-cell response (23, 55). At the same time, there are evidence that a parallel maturation of B-lineage cells might take place within the CNS: B-cell follicles are present in the meninges of patients with long-standing MS (56, 57) and could also be present already at an earlier disease stage (58). Expression of the Activation-Induced Cytidine Deaminase gene (*AICDA*), which encodes the DNA-editing deaminase involved in SHM and class switch recombination, has been detected in intrathecal B-lineage cells (59). In the present study, we detected intrathecally expanded IgM memory B cells and expanded IgM ASCs. Moreover, we found evidence of isotype switching from IgM to IgG1, from IgG1 to IgA1 and from IgG1 to IgG2, and observed extensive connections between the intrathecal memory and ASC compartments. Taken together, our data support the idea of an ongoing maturation, SHM and isotype switching of B-lineage cells within the intrathecal milieu.

Our observation of expanded and extensively mutated IgA ASCs corroborates previous studies detecting IgA in the CSF (60) and IgA-producing cells in the CSF and MS lesions (23, 61–63). A recent study found that a proportion of IgA-producing cells in the CSF recognize gut microbiota, and the authors proposed that such cells could represent a population of regulatory cells (23). In the present study, however, we did not detect any distinct regulatory phenotype of IgA-expressing B-lineage cells in the CSF. Nevertheless, our study does not rule out the presence of such regulatory cells in the brain or meninges.

This is the first study to provide a detailed clonal evolution of B-lineage cells in the CSF of MS patients based on somatic mutations in paired heavy- and light-chain sequences of single cells. In 17 out of 21 investigated patients, we were able to depict lineage trees of ASCs accurately based on mutations in full-length transcripts of both chains. In contrast to previous studies utilizing different variants of bulk amplification and sequencing of heavy-chain transcripts (27, 33, 55), we found evidence of a more focused humoral immune response characterized by smaller lineage trees with fewer offspring per ancestor node. One possible explanation would be that a bulk sequencing approach may capture a larger part of the intrathecal B-cell repertoire. On the other hand, it is also well-known that high-throughput bulk sequencing introduces PCR- and sequencing-errors, which are inherently impossible to discern from somatic mutations (64).

Single-cell studies of B-lineage cells are imperative for assessing clonal expansion and acquiring detailed knowledge of the paired V_H_:V_L_ repertoires. We successfully achieved to explore single B-lineage cells in the CSF of MS patients in an unbiased manner by combining RNA-seq technology with bioinformatics. Nevertheless, there are limitations of the study that need to be recognized. The number of ASCs in the CSF of MS patients is generally scarce, and the number of single cells sequenced and analyzed is only a small part of a larger population of B-lineage cells with a plethora of different phenotypes within the CNS. Underscoring this fact, we recently demonstrated that the clonal overlap of CSF B cells at two different time points is much lower than the overlap of the secreted CSF IgG proteome (28). This could be due to limited sampling and/or that B-lineage cells in the CSF only partly represent populations in the brain and meninges.

Our study represents one of the most detailed investigations of B-lineage cells in the CSF of MS patients to date. For the first time, we demonstrate a strong connection between a G1m allotype and the heavy- and light-chain variable gene repertoires. We anticipate that such interactions can point to biological effects of the G1m allotypes that might be of importance in humoral immune responses in MS.

## Methods

### Patients and sample collection

Twenty-four subjects with symptoms and brain magnetic resonance imaging (MRI) scans strongly suggestive of MS were recruited during diagnostic work-up at the Departments of Neurology at Akershus University Hospital and Oslo University Hospital. All patients were of Caucasian ethnicity. Three patients were excluded due to too low frequency of antibody-secreting cells and/or absence of OCBs (Supplemental Table 1). Based on MRI scans, clinical symptoms and CSF findings, 20 included patients met the criteria of relapsing-remitting MS according to the 2017 McDonald revisions (65), and one patient had developed a secondary-progressive disease (Table 1). MS19 and MS20 had been previously treated with intravenous infusions of methylprednisolone 1 g per day for three days within two weeks of inclusion. None of the other patients had received corticosteroids or any other type of immunomodulatory treatment.

CSF and serum were collected during diagnostic lumbar puncture. The first 7 ml of CSF were collected for diagnostic purposes, and 2 x 9 ml CSF were subsequently collected in 15 ml Falcon tubes. Collected CSF was centrifuged (1600 rpm, 10 min, room temperature (RT)) and cell pellets were resuspended in approximately 100 μl of supernatant. Cell count and the presence of red blood cells was checked in a Bürker counting chamber, and none of the CSF samples contained enough cells to indicate contamination of lymphocytes from blood. Serum was left at RT for at least 30 min after collection, centrifuged (2800 rpm, 10 min, RT) and stored at −80°C. It was subsequently thawed and used to determine the expression of the G1m allotypes of each subject by ELISA according to a previously published protocol (8).

### Single-cell sorting by flow cytometry

The CSF cell suspension was transferred to a V-bottom plate, centrifuged (1600 rpm, 4 min, 4°C) and the supernatant was removed. Cells were resuspended in 40 μl of LIVE/DEAD Fixable Violet (Molecular Probes L34963) according to the manufacturer’s instructions, and incubated at RT for 15 min in the dark. 10 μl of 30% bovine serum albumin (BSA) was added, followed by a mixture of the following anti-human antibodies from BD Biosciences: CD3-BV510 (HIT3a), CD14-BV510 (MφP9), CD16-BV510 (3G8), CD19-AF488 (HIB19), CD27-PE-CF594 (M-T271), CD38-APC (HIT2), IgD-APC-H7 (IA6-2). Cell suspensions were incubated for 30 min on ice, centrifuged (1600 rpm, 4 min, 4°C) and washed once with flow buffer (PBS, 0.5% BSA, 2 mM EDTA). Cells were resuspended in 200 μl of flow buffer and taken for single-cell index-sorting on a FACS ARIA III cell sorter equipped with 408, 488, 561 and 633 nm lasers (BD Biosciences) at the Flow cytometry core facility at the Oslo University Hospital. ASCs and B cells were sorted into separate plates (Figure 1). Single cells were index-sorted into 96-well plates (BioRad) containing 2 μl lysis buffer per well (1:20 RNase inhibitor (Clontech) in 0.2% (v/v) Triton X-100 (Sigma)).

### Generation of scRNA-seq libraries

Plates containing sorted single cells were spun down (2500 rpm, 5 min, 4°C) and stored at −80°C or immediately processed for cDNA synthesis. cDNA synthesis was done following the Smart-Seq2 protocol (17) with minor modifications as described below. For reverse transcription SmartScribe RT (Clontech) and corresponding buffer was used. Further, a modified TSO primer with biotinylation was introduced: Bio-AAGCAGTGGTATCAACGCAGAGTACrGrG+G. In the cDNA amplification step, ASCs underwent 21 cycles of amplification and CD19^+^ B cells were amplified with 22 cycles. Amplified cDNA was purified with 20 μl Ampure XP (Agencourt) beads per well. Tagmentation was done using in-house made Tn5 transposase (66) dually indexed with Nextera (XT) N7xx and S5xx index primers (125 nM). Purified libraries, consisting of up to 384 cells each, were sequenced on the Illumina NextSeq500 platform at the Norwegian Sequencing Centre using 75 bp paired-end read high-output sequencing runs, resulting in approximately 1 million reads per cell.

### Processing of raw sequencing data

Low-quality sequences and adapter sequences were trimmed off the raw scRNA-seq reads with Cutadapt v.1.18 (67) and Trim Galore v0.6.1 in paired-end mode. We then quantified transcript expression using Salmon v0.11.3 (68), building the salmon index using cDNA sequences from GRCh38.94 and a k-mer length of 25. Quantified transcripts were subsequently collapsed to gene level, and corrected for transcript length using tximport v1.8.0 (69). Quality control was performed in R version 3.5.3 using the scater package (70), and was based on the following measures: Number of detected genes and reads, percent mitochondrial genes, reads mapping to the reference, percent Ig genes, and successful reconstruction of at least one productively rearranged heavy chain BCR sequence by BraCeR (18). Library-specific threshold values can be found in Supplemental Table 2.

### B-cell receptor analysis

Raw reads were provided as input to BraCeR (18) in *assemble* mode in order to reconstruct full-length paired heavy and light BCR chains for each cell. We ran BraCeR assembly with --*threshold5000* for ASCs and --*threshold500* for B cells in order to filter out reconstructed BCRs likely arising from well-to-well contamination or PCR errors in indices. BraCeR yielded the full-length recombined V-region of heavy and light chains in most cells, and additionally sufficient coverage of the constant region to determine isotypes and G1m1 or G1m3 allotypes based on perfect, full-length alignment to the CH1 region of the *IGHG1*01* or *IGHG1*03* alleles, respectively. Notably, the CH1 regions of several of the G1m1-containing alleles are identical, and all cells of the G1m1 allotype were thus assigned the *IGHG1*01* allele by BraCeR.

BraCeR was subsequently used in *summarise* mode to identify clonal relationships between cells and construct lineage trees based on paired heavy and light chains. We identified clonally related B-lineage cells separately for each patient with the following parameters: --*include_multiplets --infer_lineage*. Likely cell multiplets were then manually removed as previously described (71).

Information regarding variable (V), diversity (D) and joining (J) gene segment usage, light chain, isotype, clonality etc. was extracted from the summary files generated by BraCeR and used for more extensive BCR repertoire analysis using Change-O and Alakazam v0.3.0 (72). The number of V-segment somatic mutations was identified for the most highly expressed heavy- and light chain in each cell using IMGT/HighV-QUEST (73) and normalized by the length of the V-segment of each reconstructed sequence. In order to remove ambiguous gene assignment of duplicated V genes, those gene segments were collapsed into newly created subgroups, for example *IGKV1-33* and *IGKV1D-33* were collapsed into *IGKV1(D)-33*. A Circos plot, visualizing the *IGHV4:IGKV1* gene pairing frequencies, was generated using Circos Table Viewer (74).

### IGHG1 allele inference

While the G1m allotypes expressed by each patient were determined serologically, we also computationally inferred the specific *IGHG1* alleles carried by each individual using Salmon (68) and a custom script. In short, reads from IgG1 cells were mapped to all *IGHG1* reference allele sequences obtained from IMGT, and the *IGHG1* allele with highest mapping rate to the CH1, CH2 and CH3 parts of *IGHG1* was determined as the *IGHG1* allele for each cell. Allele assignments for each cell for each patient were manually inspected, and patient-specific alleles were determined based on the most frequent allele assignments.

### Gene expression analysis and phenotyping of B-lineage cells

Subsets of B-lineage cells were initially determined according to the surface expression of CD27 and CD38 measured by flow cytometry, isotype inferred by BraCeR, and SHM load (Table 2). Ig genes are frequently expressed in B-lineage cells and were discarded from the gene expression analysis before normalization in order to avoid masking of non-Ig-related transcriptional differences. Genes with more than three reads detected in more than five cells were retained. The gene expression matrix was subset according to cells of interest, and subsequently normalized by total reads and logarithmized as X = ln(X + 1) with scanpy v1.4.4 (75). We then identified highly variable genes using the highly_variable_genes function of scanpy (*min_mean=0.1*, *max_mean=10, min_disp=0.25*). In order to remove batch effects and reduce patient-to-patient variation, we regressed out number of detected genes and reads, percent mitochondrial genes and patient-specific variation, while retaining variation explained by cell type, using NaiveDE (https://github.com/Teichlab/NaiveDE) and the highly variable genes. Lastly, the expression matrix was scaled using sklearn (scikit-learn v0.21.3) (76). We then used scanpy to run Principal Component Analysis (PCA) of the scaled expression matrices, and visualized the populations using Uniform Manifold Approximation and Projection for Dimension Reduction (UMAP) (77). The first 25 principal components were used for visualization of all the cells together, while only the first seven principal components were used for visualization of only the CD19^+^ B cells. Cell cycle phase of each cell was inferred using the score_genes_cell_cycle function of scanpy using the provided regev_lab_cell_cycle_genes.txt file for specifying genes associated with the S and G2M phases.

### Statistics

Statistical analysis and plots were made using R version 3.5.3 and ggpubr v0.2.4. SHM frequencies were compared between all sequences of different isotypes or B-lineage populations using a Wilcoxon rank-sum test, while median SHM frequencies for each patient were compared across B-lineage populations with a Wilcoxon signed-rank test. Frequencies of G1m1 cells were compared within ASCs and memory B cells using a binomial test. Paired t-test was used to analyze differences in G1m1 usage between ASCs and memory B cells and between G1m1 and G1m3 cells in G1m1/G1m3 heterozygous patients. Wilcoxon rank-sum test was used to test differences in Igκ usage for IgG1 ASCs expressing G1m1 or G1m3 and proportion of cells using *IGHV* family genes depending on the expressed G1m allotype. All tests were two-sided with a significance level of 0.05 unless otherwise stated in the figure legends.

### Data deposition

The raw scRNA-seq fastq files, reconstructed BCR sequences and accompanying metadata have been deposited at the European Genome-Phenome Archive (EGA) under accession number XXXXXX, with controlled access.

### Study approval

The study was approved by the Committee for Research Ethics at the South-Eastern Norwegian Health Authority (2009/23S-04143a). All patients signed a written informed consent.

## Author contributions

AL, JP, LMS, FV, and TH conceived the study and designed the experiments. AL, RH, TH, and PBH recruited patients and provided patient data. AL, IL, JP, SWQ, and FV conducted experiments. IL, JP and AL conducted data analysis. AL, IL, and JP drafted the manuscript. All authors contributed in revising the manuscript.

## Acknowledgements

We would like to express our gratitude to patients who participated in this study. We thank Łukasz Wyrożemski for invaluable assistance in performing experiments. The study was funded by grants from the Norwegian Women’s Public Health Association, the University of Oslo World-leading research program on human immunology (WL-IMMUNOLOGY), and by Stiftelsen KG Jebsen (project SKGJ-MED-017). In addition, A.L. is the recipient of Per B. Larsen’s grant 2018, Odd Fellow’s research grant 2019, and the Norwegian Neurological Association research prize in multiple sclerosis 2019 provided by Sanofi Genzyme.

