## Supplemental Materials for "Stereotyped B-cell responses are linked to IgG constant region polymorphisms in multiple sclerosis"

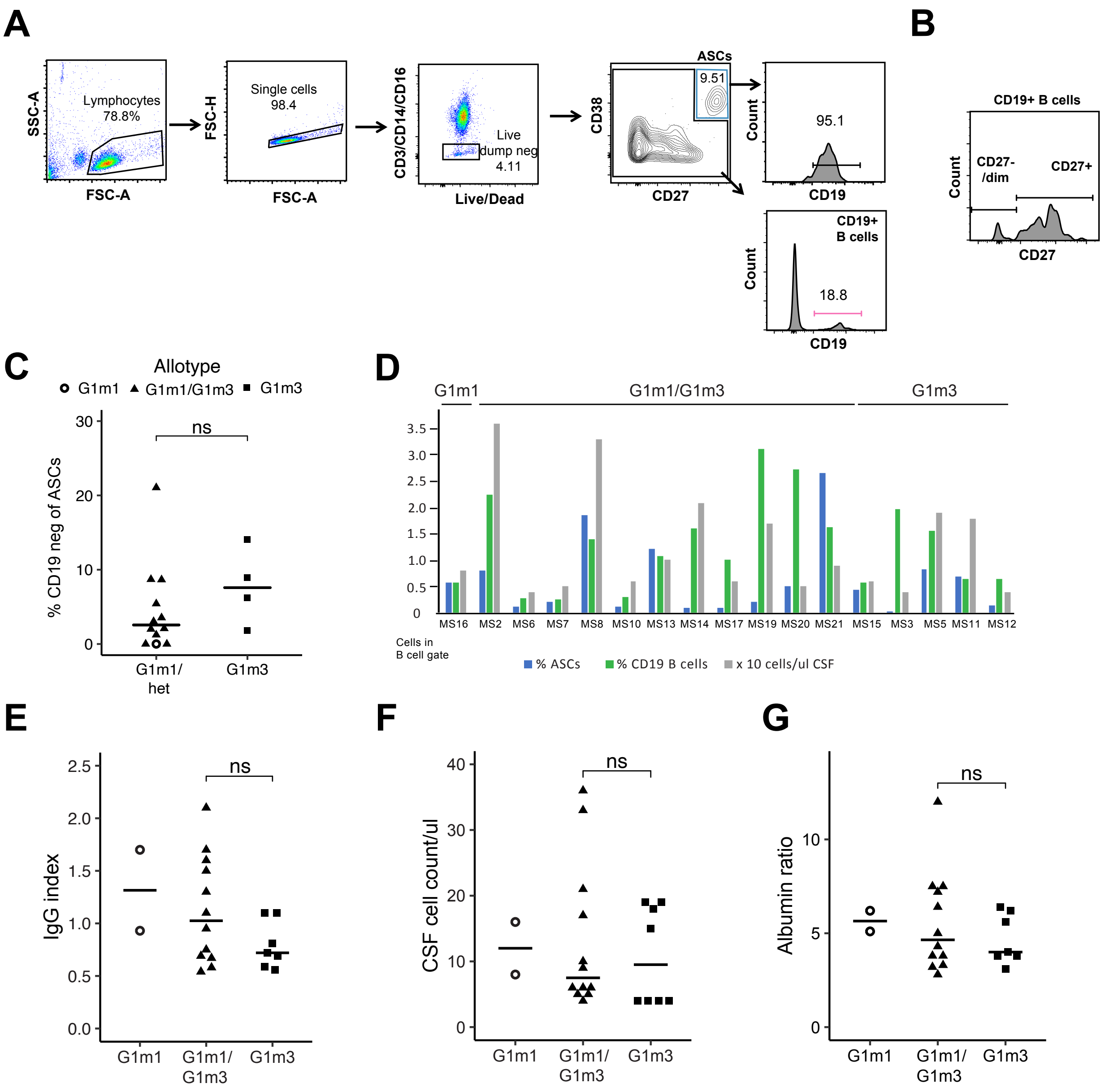


**Figure S1. Analysis of intrathecal B-lineage cells and related parameters stratified by G1m allotype. A.** Representative flow cytometry gating strategy for one patient showing sorting and analysis gates. **B.** Representative flow cytometry gating strategy for one patient showing gating of index-sorted CD19$^{+}$ B cells into CD27$^{+}$ and CD27$^{-/dim}$ populations. **C.** Percent of ASCs being CD19 negative according to flow cytometry analysis. Data for all patients with a substantial number of recorded events are summarized (n=17). Differences between patient groups were tested using an unpaired t-test. **D.** Percent ASCs and CD19$^{+}$ B cells according to flow cytometry analysis for all patients with a substantial number of recorded events (n=17). CSF cell count is displayed in gray. **E,F,G.** IgG index (E), CSF cell count (F) and albumin ratio (G) for included patients, stratified by patient allotype. Differences between G1m1/G1m3 heterozygotes and G1m3 homozygotes were tested using unpaired t-tests. Ns = p>0.05.

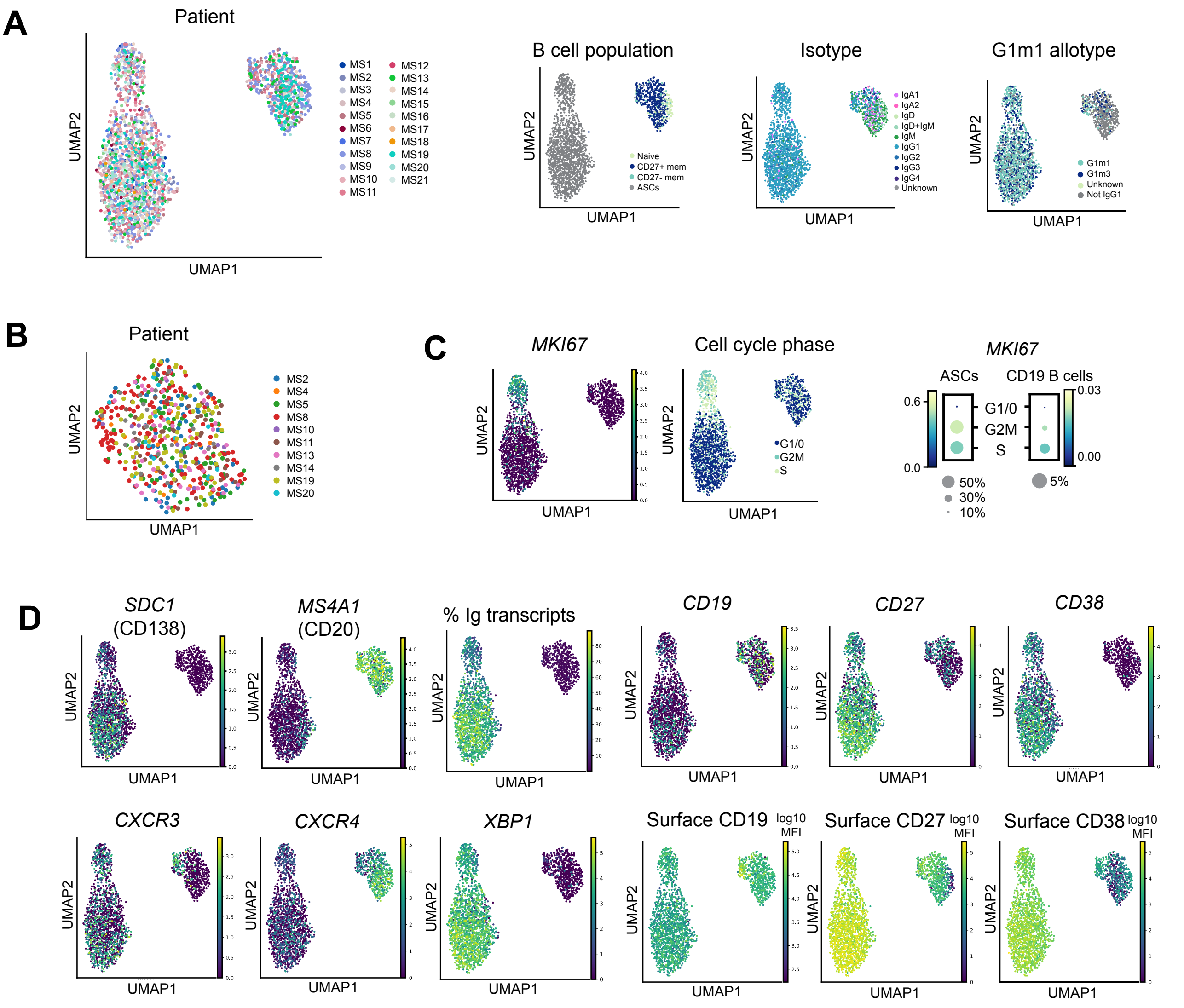


**Figure S2. Transcriptional profiling of cerebrospinal B-lineage cells in MS.** The transcriptional profile of B-lineage cells from the CSF of 21 MS patients were visualized using UMAP as in Figure 2B. **A.** UMAP projection of all B-lineage cells (2165 cells) from all 21 patients, colored by patient, B cell population, isotype and G1m1 allotype. **B.** UMAP projection of all $CD19^{+}$ B-lineage cells (544 cells) from all 10 patients for which this cell population was sorted, colored by patient. **C.** UMAP projections as in (A), colored by *MKI67* expression (left) and inferred cell cycle (right) without using *MKI67* in the list of proliferation-associated genes. Dotplots show expression of *MKI67* in ASCs and $CD19^{+}$ B cells according to the cell cycle phases shown in the UMAP plot. **D.** UMAP projections as in (A), colored by genes of interest or median fluorescence intensity (MFI) of the cell surface markers CD19, CD27 and CD38 (bottom row) obtained during index-sorting.

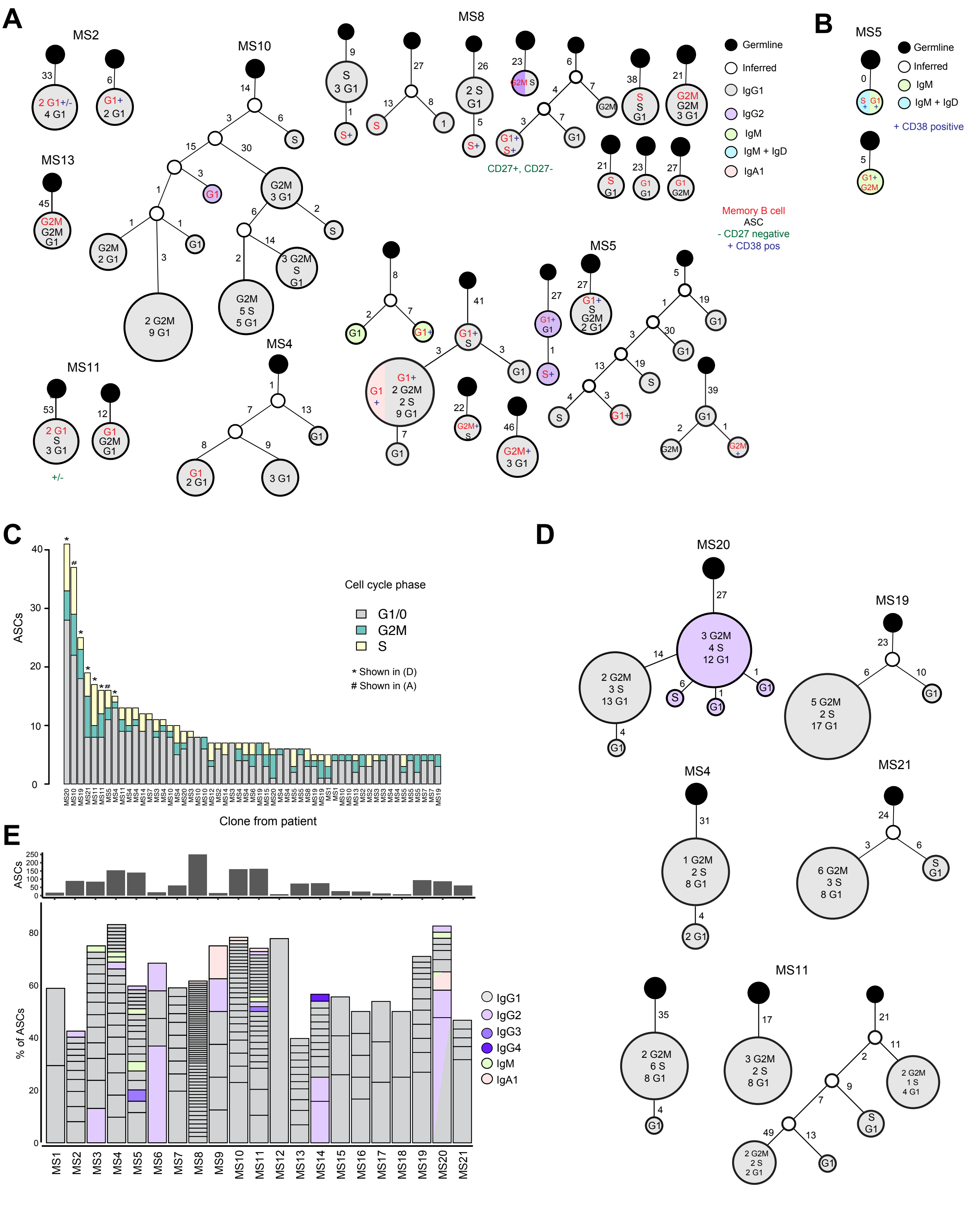


**Figure S3. Clonal connections across cell cycle stages and isotypes. A.** Lineage trees consisting of ASCs and memory B cells with annotated cell cycle stage and CD38 and CD27 cell surface expression. **B.** Lineage trees restricted to memory B cells with annotated cell cycle stage and surface expression of CD38 and CD27. **C.** Cell cycle stage of largest expanded clone groups. **D.** Lineage trees for largest expanded clone groups. **E.** Isotype of clone groups determined by BraCeR.

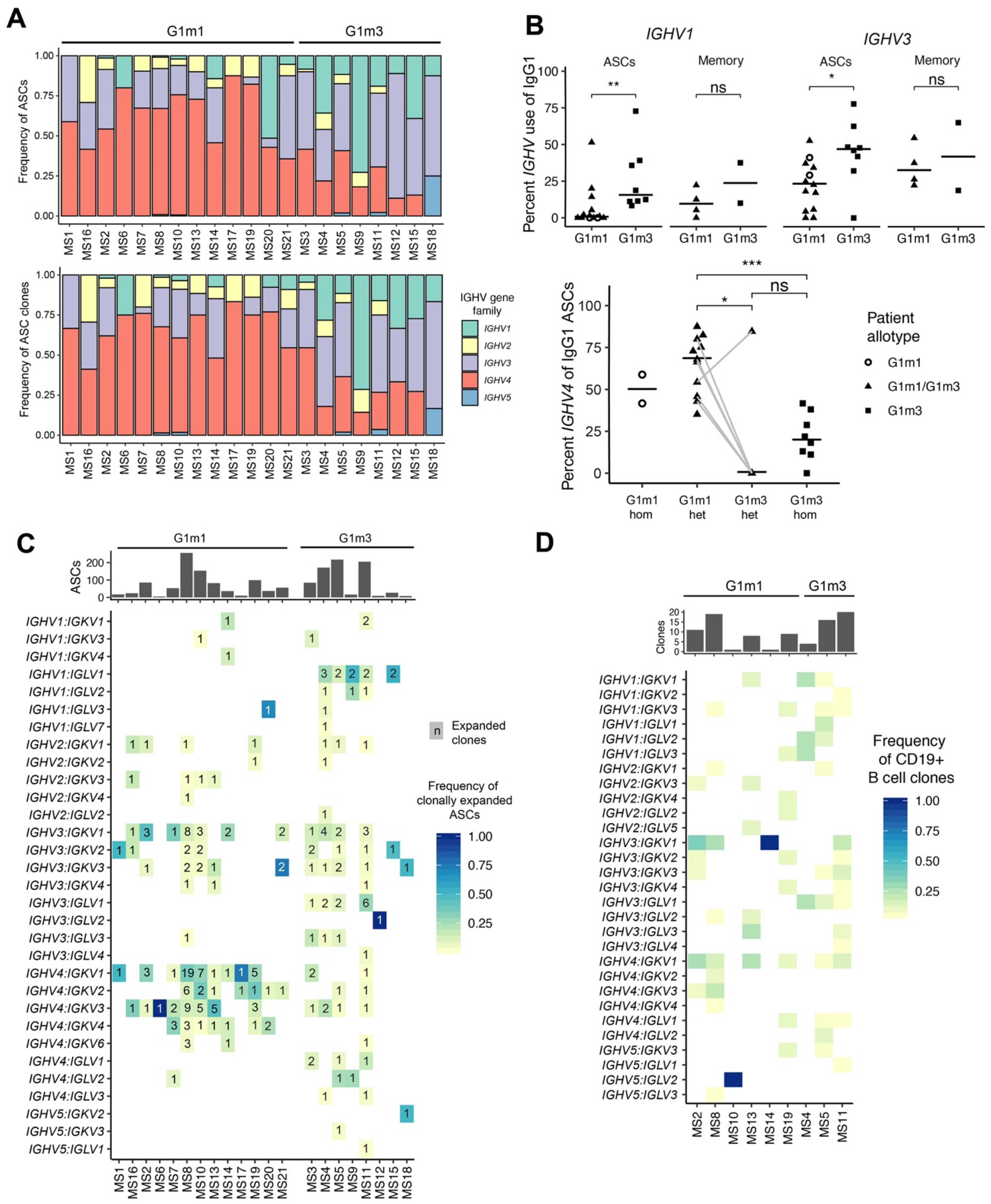


**Figure S4. *IGHV* gene usage in B-lineage cells from the cerebrospinal fluid. A.** *IGHV* gene family usage in ASCs in each patient, grouped by G1m1 carrier status. **B.** *IGHV1* and *IGHV3* gene family usage in G1m1 and G1m3 ASCs and memory B cells (upper panel). Lower panel shows *IGHV4* gene family usage in ASCs for G1m1 cells coming from homozygous or heterozygous patients and G1m3 cells from homozygous or heterozygous patients. Each dot represents a patient; horizontal line is median for all patients in the given group. Lines connect G1m1 and G1m3 ASCs from the same patient. Two of the heterozygous patients had no ASCs expressing G1m3. Statistical differences were tested using paired t-test – G1m1 versus G1m3 cells from heterozygotes and Wilcoxon rank-sum test for unpaired data (G1m1 cells from heterozygous patients versus G1m3 cells from homozygous. **C.** Heatmap showing $V_H$:$V_L$ gene segment pairing frequencies for clonally expanded G1m1 and G1m3 ASCs only. The numbers within each square indicate the number of clone groups using each gene combination, while the color represents frequency of total number of clonally expanded ASCs. **D.** Heatmap showing $V_H$:$V_L$ gene segment pairing frequencies for G1m1 and G1m3 memory B cells, with cells belonging to a clone group being treated as one.

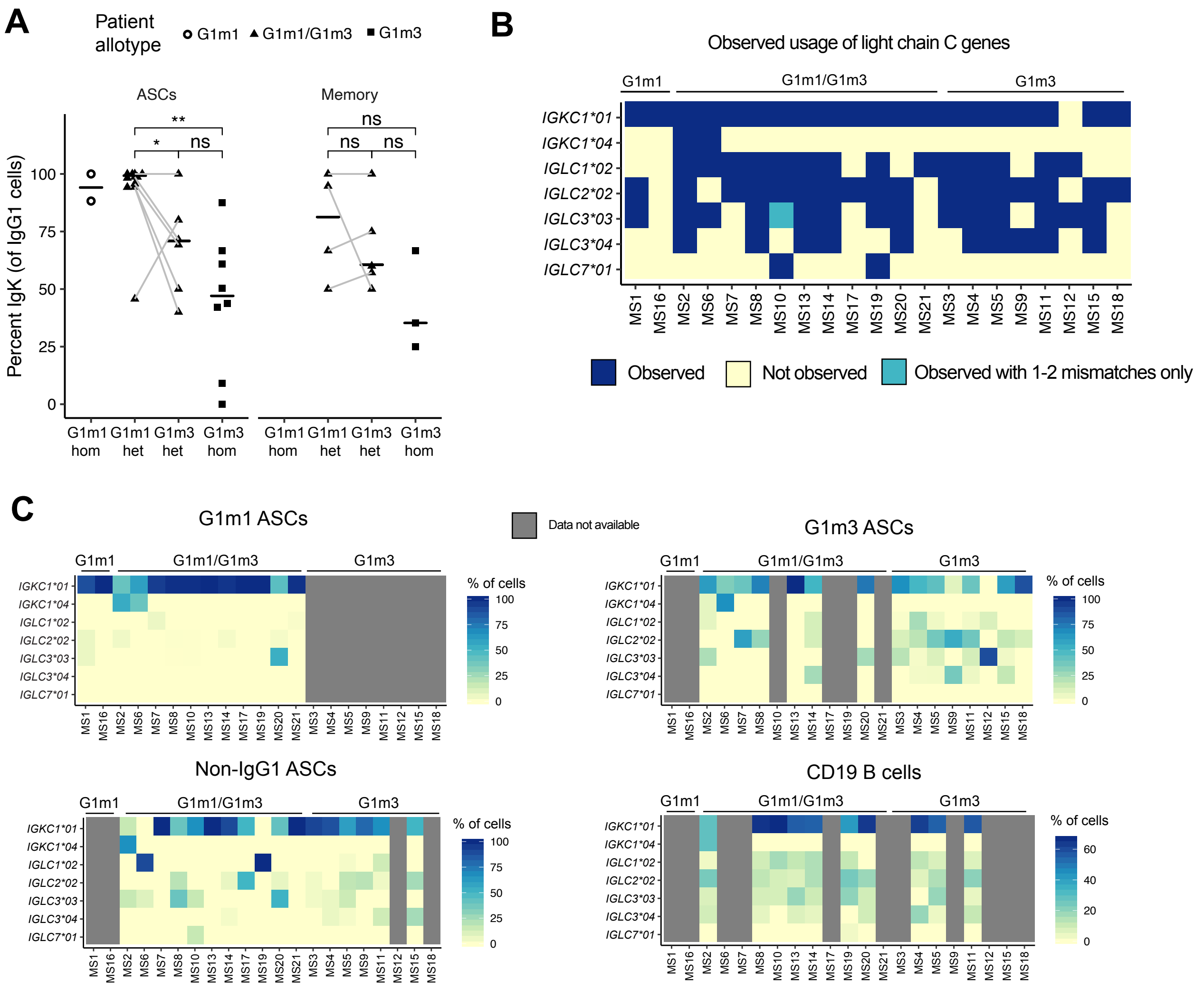


**Figure S5. Light chain constant gene usage in B-lineage cells from the cerebrospinal fluid. A.** Frequency of ASCs and memory B cells using κ light chain for each patient, grouped by patient allotype. Each dot represents a patient; cell populations from the same patients are connected by a line. Two of the heterozygous patients had no G1m3 ASCs. Statistical differences between G1m1 and G1m3 cells from heterozygotes were tested using a paired t-test. Differences between G1m1 cells from heterozygous patients and G1m3 cells from homozygous patients were compared by Wilcoxon rank-sum test. **B.** Observed usage (presence or absence in at least one B-lineage cell) of specific Ig light chain constant region genes and alleles in each patient. Constant region reconstructed by BraCeR was compared to germline reference sequences to determine perfect, full-length matches to each allele. **C.** Frequency of usage of specific Ig light chain constant region genes and alleles in each patient, stratified by allotype, isotype and cell population.

Table S1. Excluded patients.

| ID | Sex | Age | Diagnosis | Disease duration (months) | Rela pses | Months since last relapse | CSF cell count$^{A}$ | OCB$^{B}$ | Album in ratio | IgG index | Allotype$^{C}$ | Reason for exclusion |
|---|---|---|---|---|---|---|---|---|---|---|---|---|
| **MS22** | F | 42 | RR-MS | 12 | 1 | 12 | 4 | + | 6.6 | 0.64 | G1m3 | Few sorted cells (8 ASCs) |
| **MS23** | M | 27 | RR-MS | 2 | 1 | 2 | 4 | - | 9.0 | 0.53 | G1m1/G1 m3 | No OCBs; few sorted cells (2 ASCs) |
| **MS24** | F | 38 | Clinically isolated syndrome | 168 | 1 | 168 | 4 | - | 6.6 | 0.58 | G1m1 | No OCBs; few sorted cells (2 ASCs) |

F-female; M-male; RR-MS – relapsing-remitting multiple sclerosis; ND – not determined.

[A]mononuclear cells count per microliter of CSF

[B]positive (+) or negative (-) for more than 2 CSF restricted oligoclonal bands on isoelectric focusing

[C]determined in serum

Table S2. Library-specific threshold values for quality control (QC) of scRNA-seq data.

| **Library** | **Subjects** | **Cell type** | **Min. reads** | **Max. reads** | **Min. genes** | **Max. genes** | **% mapped reads** | **% Ig reads** | **% mt reads** | **BCR** | **Total cells** | **QC pass** |
|---|---|---|---|---|---|---|---|---|---|---|---|---|
| **scLib1** | MS1 | ASC | 1x$10^5$ | 2.5x$10^6$ | 1000 | 5800 | >40 | >10 | <8 | Productive IgH | 29 | 17 |
| | MS2 | B | | | | | | <10 | | | 93 | 71 |
| | | ASC | | | | | | >10 | | | 96 | 89 |
| | MS3 | ASC | | | | | | >10 | | | 96 | 84 |
| **scLib2** | MS4 | B | 1x$10^5$ | 1.2x$10^6$ | 1200 | 5100 | >40 | <10 | <8 | Productive IgH | 96 | 17 |
| | | ASC | | | | | | >10 | | | 192 | 154 |
| | MS7 | ASC | | | | | | | | | 77 | 61 |
| | MS9 | ASC | | | | | | | | | 19 | 16 |
| **scLib3** | MS5 | B | 1x$10^5$ | 1.8x$10^6$ | 1000 | 6100 | >40 | <10 | <8 | Productive IgH | 90 | 81 |
| | | ASC | | | | | | >10 | | | 167 | 139 |
| | MS6 | ASC | | | | | | | | | 22 | 19 |
| | MS8 | ASC | | | | | | | | | 96 | 79 |
| **scLib4** | MS8 | B | 1.5x$10^5$ | 1.9x$10^6$ | 1000 | 6200 | >40 | <10 | <8 | Productive IgH | 192 | 159 |
| | | ASC | | | | | | >10 | | | 192 | 171 |
| | MS11 | B | | | | | | <10 | | | 80 | 45 |
| | | ASC | | | | | | >10 | | | 192 | 162 |
| **scLib6** | MS10 | B | 1.5x$10^5$ | 2x$10^6$ | 1000 | 5800 | >40 | <10 | <8 | Productive IgH | 70 | 12 |
| | | ASC | | | | | | >10 | | | 208 | 161 |
| **scLib7** | MS12 | ASC | 1.5x$10^5$ | 2.7x$10^6$ | 1000 | 6000 | >40 | >10 | <8 | Productive IgH | 14 | 9 |
| | MS13 | B | | | | | | <10 | | | 96 | 47 |
| | | ASC | | | | | | >10 | | | 103 | 73 |
| | MS14 | B | | | | | | <10 | | | 44 | 22 |
| | | ASC | | | | | | >10 | | | 96 | 76 |
| **scLib8** | MS16 | ASC | 1.5x$10^5$ | 2.7x$10^6$ | 800 | 6000 | >40 | >10 | <8 | Productive IgH | 34 | 24 |
| | MS20 | ASC | | | | | | >10 | | | 96 | 86 |
| | | B | | | | | | <10 | | | 96 | 16 |
| **scLib9** | MS15 | ASC | 1.5x$10^5$ | 2.7x$10^6$ | 1000 | 6000 | >40 | >10 | <8 | Productive IgH | 31 | 27 |
| | MS17 | ASC | | | | | | | | | 18 | 13 |
| | MS18 | ASC | | | | | | | | | 8 | 8 |
| | MS19 | ASC | | | | | | | | | 96 | 93 |
| | | B | | | | | | <10 | | | 112 | 74 |
| | MS21 | ASC | | | | | | >10 | | | 88 | 60 |
| **Total** | | | | | | | | | | | **3052** | **2165** |

IgH = immunoglobulin heavy chain, mt = mitochondrial.

Table S3. Number of analyzed cells from CSF B-cell populations.

| Subject ID | B-cell subsets | | | | | | |
|---|---|---|---|---|---|---|---|
| | | | Memory | | | | |
| | Antibody secreting cells | Naïve | Non-switched CD27$^{-/dim}$ | Non-switched CD27$^{+}$ | Switched CD27$^{-/dim}$ | Switched CD27$^{+}$ | Unclassified |
| **MS1** | 17 | - | - | - | - | - | - |
| **MS2** | 89 | 3 | 3 | 24 | 3 | 38 | - |
| **MS3** | 84 | - | - | - | - | - | - |
| **MS4** | 154 | 4 | 1 | 5 | 0 | 7 | - |
| **MS5** | 139 | 4 | 2 | 33 | 1 | 41 | - |
| **MS6** | 19 | - | - | - | - | - | - |
| **MS7** | 61 | - | - | - | - | - | - |
| **MS8** | 250 | 55 | 6 | 49 | 7 | 40 | 2 |
| **MS9** | 16 | - | - | - | - | - | - |
| **MS10** | 161 | 2 | - | 3 | 1 | 6 | - |
| **MS11** | 162 | 2 | - | 14 | 2 | 27 | - |
| **MS12** | 9 | - | - | - | - | - | - |
| **MS13** | 73 | 1 | 2 | 18 | 2 | 24 | - |
| **MS14** | 76 | 5 | 2 | 5 | - | 9 | 1 |
| **MS15** | 27 | - | - | - | - | - | - |
| **MS16** | 24 | - | - | - | - | - | - |
| **MS17** | 13 | - | - | - | - | - | - |
| **MS18** | 8 | - | - | - | - | - | - |
| **MS19** | 93 | 8 | 3 | 17 | 3 | 43 | - |
| **MS20** | 86 | 1 | 2 | 9 | 0 | 4 | |
| **MS21** | 60 | - | - | - | - | - | - |
| **Total** | **1621** | **85** | **21** | **177** | **19** | **239** | **3** |

Table S4. Clonal expansion of antibody secreting cells

| Patient ID | % clonal cells | Size of clonotypes | Expanded clonotypes | Lineage tracing | | | | |
|---|---|---|---|---|---|---|---|---|
| | | | | Lineage trees | Unique sequences per tree | Clonotype size | SHM (range) within tree | Isotype |
| MS1 | 58.8 | 5 | 2 | 0 | - | - | - | - |
| MS2 | 42.5 | 2-7 | 12 | 1 | 2 | 5 | 44-50 | IgG1 |
| MS3 | 75.0 | 2-11 | 14 | 5 | 3 | 9 | 35-49 | IgG1 |
| | | | | | 2 | 3 | 27-31 | IgG1 |
| | | | | | 3 | 5 | 23-25 | IgG1 |
| | | | | | 2 | 11 | 47-50 | IgG2 |
| | | | | | 2 | 4 | 19-22 | IgG1 |
| MS4 | 83.1 | 2-15 | 23 | 3 | 3 | 6 | 14-17 | IgG1 |
| | | | | | 2 | 2 | 23-26 | IgG1 |
| | | | | | 2 | 13 | 31-35 | IgG1 |
| MS5 | 59.7 | 2-16 | 20 | 7 | 4 | 16 | 44-51 | IgG1 |
| | | | | | 3 | 5 | 12-22 | IgM |
| | | | | | 3 | 3 | 47-58 | IgG1 |
| | | | | | 4 | 4 | 24-37 | IgG1 |
| | | | | | 3 | 6 | 56-59 | IgG1 |
| | | | | | 2 | 2 | 39-41 | IgG1 |
| | | | | | 4 | 6 | 42-46 | IgG3 |
| MS6 | 68.4 | 2-7 | 4 | 2 | 2 | 2 | 50-53 | IgG1 |
| | | | | | 2 | 2 | 36-41 | IgG2 |
| MS7 | 65.6 | 2-12 | 10 | 4 | 2 | 2 | 32-33 | IgG1 |
| | | | | | 2 | 3 | 68-75 | IgG1 |
| | | | | | 3 | 5 | 29-33 | IgG1 |
| | | | | | 2 | 5 | 36-40 | IgG1 |
| MS8 | 61.6 | 2-6 | 61 | 28 | 2 | 2 | 32-34 | IgG1 |
| | | | | | 2 | 2 | 21-32 | IgG1 |
| | | | | | 2 | 2 | 25-45 | IgG1 |
| | | | | | 2 | 3 | 22-23 | IgG1 |
| | | | | | 2 | 2 | 31-32 | IgG1 |
| | | | | | 3 | 4 | 4-15 | IgG1 |
| | | | | | 2 | 1 | 27-34 | IgG1 |
| | | | | | 2 | 2 | 26-31 | IgG1 |
| | | | | | 2 | 2 | 29-30 | IgG1 |
| | | | | | 2 | 2 | 23-24 | IgG1 |
| | | | | | 2 | 2 | 18 | IgG1 |
| | | | | | 2 | 3 | 20-24 | IgG1 |
| | | | | | 2 | 2 | 17-18 | IgG1 |
| | | | | | 2 | 2 | 24-27 | IgG1 |
| | | | | | 2 | 2 | 29-32 | IgG1 |
| | | | | | 2 | 2 | 36-37 | IgG1 |
| | | | | | 2 | 2 | 18-22 | IgG1,IgM |
| | | | | | 3 | 3 | 34-41 | IgG1 |
| | | | | | 2 | 2 | 26-28 | IgG1 |
| | | | | | 2 | 2 | 27-28 | IgG1 |
| | | | | | 2 | 2 | 15-21 | IgG1 |
| | | | | | 2 | 2 | 39-42 | IgG1 |
| | | | | | 2 | 2 | 14-20 | IgG1 |
| | | | | | 2 | 3 | 28-29 | IgG1 |
| | | | | | 2 | 2 | 11-19 | IgG1 |
| | | | | | 2 | 4 | 22-27 | IgG1 |
| | | | | | 2 | 2 | 16-20 | IgG1 |
| | | | | | 2 | 2 | 19 | IgG1 |

| | | | | | | | | |
|---|---|---|---|---|---|---|---|---|
| MS9 | 75.0 | 2 | 6 | 2 | 2 | 2 | 52 | IgA1 |
| | | | | | 2 | 2 | 21-29 | IgG1 |
| MS10 | 78.3 | 2-37 | 26 | 16 | 8 | 37 | 20-67 | IgG1 |
| | | | | | 3 | 4 | 19-28 | IgG1 |
| | | | | | 3 | 4 | 26-38 | IgG1 |
| | | | | | 2 | 2 | 33-65 | IgA1 |
| | | | | | 5 | 5 | 15-21 | IgG1 |
| | | | | | 2 | 2 | 20-27 | IgG1 |
| | | | | | 2 | 2 | 74-94 | IgA1 |
| | | | | | 5 | 8 | 33-39 | IgG1 |
| | | | | | 2 | 2 | 14-15 | IgG1 |
| | | | | | 3 | 3 | 19-21 | IgG1 |
| | | | | | 2 | 2 | 23-28 | IgG1 |
| | | | | | 2 | 8 | 26-29 | IgG1 |
| | | | | | 2 | 10 | 33-34 | IgG1 |
| | | | | | 2 | 3 | 19-21 | IgG1 |
| | | | | | 2 | 2 | 26-31 | IgG1 |
| | | | | | 2 | 2 | 40-46 | IgG1 |
| MS11 | 74.1 | 2-17 | 31 | 11 | 2 | 2 | 28-32 | IgG1 |
| | | | | | 4 | 16 | 32-79 | IgG1 |
| | | | | | 2 | 4 | 38-45 | IgG1 |
| | | | | | 2 | 2 | 46-49 | IgG1 |
| | | | | | 2 | 2 | 27-28 | IgG1 |
| | | | | | 2 | 2 | 32 | IgG1 |
| | | | | | 2 | 3 | 22-32 | IgG1 |
| | | | | | 2 | 3 | 22-27 | IgG1 |
| | | | | | 2 | 17 | 35-39 | IgG1 |
| | | | | | 2 | 2 | 59-60 | IgA1 |
| | | | | | 2 | 2 | 18-19 | IgG1 |
| MS12 | 77.8 | 7 | 1 | 0 | - | - | - | - |
| MS13 | 39.7 | 2-5 | 11 | 3 | 2 | 5 | 11-27 | IgG1 |
| | | | | | 2 | 4 | 23-24 | IgG1 |
| | | | | | 2 | 3 | 42-46 | IgG1 |
| MS14 | 56.6 | 2-12 | 13 | 3 | 2 | 2 | 36-44 | IgG1 |
| | | | | | 2 | 2 | 41-51 | IgG1 |
| | | | | | 2 | 12 | 19-24 | IgG2 |
| MS15 | 55.6 | 4-7 | 3 | 2 | 2 | 4 | 30-31 | IgG1 |
| | | | | | 4 | 7 | 25-30 | IgG1 |
| MS16 | 50.0 | 2-4 | 5 | 0 | - | - | - | - |
| MS17 | 53.8 | 2-3 | 3 | 1 | 2 | 2 | 18 | IgG1 |
| MS18 | 50.0 | 2 | 2 | 0 | - | - | - | - |
| MS19 | 71.0 | 2-25 | 12 | 5 | 2 | 5 | 21-25 | IgG1 |
| | | | | | 2 | 25 | 29-33 | IgG1 |
| | | | | | 3 | 4 | 39-48 | IgG1 |
| | | | | | 2 | 6 | 29-32 | IgG1 |
| | | | | | 2 | 4 | 33 | IgG1 |
| MS20 | 82.6 | 2-41 | 9 | 3 | 6 | 41 | 27-45 | IgG1,IgG2 |
| | | | | | 3 | 6 | 16-19 | IgA1,IgM |
| | | | | | 2 | 4 | 27-33 | IgG1 |
| MS21 | 46.7 | 2-19 | 5 | 2 | 2 | 2 | 31-35 | IgG1 |
| | | | | | 2 | 19 | 27-30 | IgG1 |

Percent clonal cells refers to the ASCs population. Number of cells detected in the smallest and largest expanded clone groups are specified as size of clonotypes. Lineage trees were created for clones comprising cells with different mutation patterns. SHM was calculated as number of mutations in heavy and light chain compared to inferred IMGT germline sequences.
